# Placental DNA methylation signatures of maternal smoking during pregnancy and potential impacts on fetal growth

**DOI:** 10.1101/663567

**Authors:** Todd M. Everson, Marta Vives-Usano, Emie Seyve, Andres Cardenas, Marina Lacasaña, Jeffrey M. Craig, Corina Lesseur, Emily R. Baker, Nora Fernandez-Jimenez, Barbara Heude, Patrice Perron, Beatriz Gónzalez-Alzaga, Jane Halliday, Maya A. Deyssenroth, Margaret R. Karagas, Carmen Íñiguez, Luigi Bouchard, Pedro Carmona-Sáez, Yuk J. Loke, Ke Hao, Thalia Belmonte, Marie A. Charles, Jordi Martorell-Marugán, Evelyne Muggli, Jia Chen, Mariana F. Fernández, Jorg Tost, Antonio Gómez-Martín, Stephanie J. London, Jordi Sunyer, Carmen J. Marsit, Johanna Lepeule, Marie-France Hivert, Mariona Bustamante

## Abstract

Maternal smoking during pregnancy (MSDP) contributes to poor birth outcomes, in part through disrupted placental functions, which may be reflected in the placental epigenome. We meta-analyzed the associations between MSDP and placental DNA methylation (DNAm) and between DNAm and birth outcomes within the Pregnancy And Childhood Epigenetics (PACE) consortium (7 studies, N=1700, 344 with any MSDP). We identified 1224 CpGs that were associated with MSDP, of which 341 associated with birth outcomes and 141 associated with gene expression. Only 6 of these CpGs were consistent with the findings from a prior meta-analysis of cord blood DNAm, demonstrating substantial tissue-specific responses to MSDP. The placental MSDP associated CpGs were enriched for growth-factor signaling, hormone activity, inflammation, and vascularization, which play important roles in placental function. We demonstrate links between placental DNAm, MSDP and poor birth outcomes, which may better inform the mechanisms through which MSDP impacts placental function and fetal growth.

---

Almost 1 in 10 pregnancies are impacted by the effects of maternal smoking during pregnancy (MSDP), with state-specific prevalence ranging from as low as 1.8% to as high as 27.1% in the USA^1^, while in Europe, the prevalence of MSDP is estimated to range between 4.2% and 18.9%^2^. Consequently, the numerous health effects of MSDP remain a significant public health concern. The impact of this exposure on fetal development has been the source of significant investigation, resulting in MSDP being recognized as a cause of multiple negative pregnancy and birth outcomes^3^.

The mechanisms that underlie this reproductive and developmental toxicity are partially understood and include molecular and anatomical changes of the placenta^4,5^. Additionally, experimental mouse models have recently highlighted the critical roles of proper placental function in ensuring successful pregnancy outcomes^6^. Epigenetic responses to prenatal exposures have emerged as potential intermediate links between early life exposures and developmental health outcomes, and epidemiologic studies of DNAm, particularly multi-cohort collaborative efforts, are powerful approaches to investigate these types of research questions^7^. So far, most studies of MSDP and epigenetics have focused on DNA methylation (DNAm) variations in cord blood, though some studies of placenta, peripheral blood and lung tissues have also been performed^8^. The Pregnancy and Childhood Epigenetics (PACE) consortium^9^ published a large meta-analysis identifying thousands of MSDP-associated variations in DNAm within cord blood and child peripheral blood^10^. However, the placental epigenome has not been as thoroughly studied, though the placenta is likely a critical target organ of MSDP-associated toxicity. A handful of prior studies have examined the relationships between MSDP and (DNAm) in human placenta^11–14^, identifying MSDP-associated loci some of which have been suggested partially mediate the effects of MSDP on lower birth weight^15^.

These studies have begun to characterize the impact that MSDP has on the human placental epigenome but have been limited by small sample sizes and have not adjusted for placental tissue heterogeneity. We aimed to address these gaps by performing a fixed effects meta-analysis examining the relationships between MSDP and variations in the placental methylome across seven independent studies that are members of the PACE consortium. We also aimed to gain insights into the potential biological and health-related impacts of these associations by performing additional analyses with nearby mRNA expression, as well as functional, regulatory, and phenotypic enrichment, and a secondary meta-analysis of the associations between placental DNAm and birth outcomes. This study represents the largest and most comprehensive examination of MSDP associations with placental DNAm in human populations and provides novel insights into the placental molecular responses to MSDP and how these relate to fetal development.

## Results

### Study population

Seven American, Australian, and European studies (N=1,700) contributed to the epigenome-wide association study (EWAS) linking MSDP to placental DNAm: including Asking Questions about Alcohol in pregnancy (AQUA)^16^, Study on the pre- and early postnatal determinants of child health and development (EDEN)^17^, Genetics of Glucose regulation in Gestation and Growth (Gen3G)^18^, Genetics, Early Life Environmental Exposures and Infant Development in Andalusia (GENEIDA), Environment and Childhood Project (INMA)^19^, New Hampshire Birth Cohort Study (NHBCS)^20^ and Rhode Island Child Health Study (RICHS)^21^. For this meta-analysis, 344 (20.2%) mothers reported any MSDP, defined as any cigarette smoking during any trimester of pregnancy. Any MSDP tended to be less prevalent in the cohorts from Canada and the USA compared to those from Australia and Europe (Table 1). Three cohorts (N=795, EDEN, GENEIDA and INMA) contributed to the EWAS of sustained MSDP, defined as maternal smoking throughout pregnancy, among which 163 (20.5%) mothers reported sustained MSDP. Distributions of covariates by cohort are provided in the supplementary materials (Excel Table S1).

**Table 1:** Distribution of any and sustained MSDP within participating cohorts.

| Cohort | Country | Any MSDP |  |  | Sustained MSDP |  |  |
| --- | --- | --- | --- | --- | --- | --- | --- |
|  |  | Total N | N (%) Non-smokers | N (%) Smokers | Total N | N (%) Non-smokers | N (%) Smokers |
| AQUA | Australia | 99 | 75 (75.76%) | 24 (24.24%) | - | - | - |
| EDEN | France | 647 | 446 (68.93%) | 201 (31.07%) | 570 | 446 (68.93%) | 124 (21.75%) |
| Gen3G | Canada | 151 | 138 (91.39%) | 13 (8.61%) | - | - | - |
| GENEID A | Spain | 87 | 67 (77.01%) | 20 (22.99%) | 82 | 67 (77.01%) | 15 (18.29%) |
| INMA | Spain | 166 | 119 (71.69%) | 47 (28.31%) | 143 | 119 (71.69%) | 24 (16.78%) |
| NHBCS | USA | 310 | 290 (93.55%) | 20 (6.45%) | - | - | - |
| RICHs | USA | 240 | 221 (92.08%) | 19 (7.92%) | - | - | - |
| TOTAL |  | 1700 | 1356 (79.76%) | 344 (20.24%) | 795 | 632 (79.50%) | 163 (20.50%) |

MSDP = Maternal smoking during pregnancy.

### Genome-wide DNAm meta-analyses

We produced four statistical models for each CpG site, modeling the associations between DNAm with both any and sustained MSDP, with and without adjustment for putative cellular heterogeneity which was estimated with a reference free deconvolution algorithm^22^. Models were adjusted for maternal age, parity, and maternal education. Genomic inflation factors from the meta-analyses ranged from λ=2.0 to 2.8 (Excel Table S2, Supplemental Figure S1). Any MSDP was associated with DNAm at 532 CpG sites after Bonferroni adjustment (Excel Table S3), while sustained MSDP was associated with DNAm at 568 CpG sites (Excel Table S4). After adjusting for putative cellular heterogeneity, any MSDP was associated with 878 CpGs (Excel Table S5), while sustained MSDP was associated with 894 CpGs (Excel Table S6). Among these, 10.7% of models for any MSDP and 3.9% of models for sustained MSDP, not adjusted for cell type, yielded heterogeneity p-values < 0.01, while only 5.5% of the models for any MSDP and 3.5% for sustained MSDP, adjusted for putative cellular heterogeneity, produced heterogeneity p-values < 0.01. Thus, heterogeneity in the associations across cohorts was lowest in the analyses that were adjusted for putative cellular heterogeneity and the CpGs from these models that yielded Bonferroni-significant associations (1224 unique CpGs) were carried forward for secondary analyses.

Overall, the Bonferroni significant CpGs were distributed throughout the genome, and the majority (68%) of CpGs exhibited lower DNAm in association with any and sustained MSDP (Figure 1). Of the 548 CpGs that yielded Bonferroni significant associations with both any and sustained MSDP, the absolute values of the parameter estimates were greater for the models of sustained MSDP compared to the estimates for any MSDP (Excel Table S7). In a secondary analysis that restricted the any MSDP models to the same three cohorts that contributed to the sustained MSDP models (EDEN, GENEIDA, and INMA), we again found that almost all CpGs (544 out of 548) exhibited increased magnitudes of association with increased duration of exposure (Excel Table S7, Supplemental Figure S2).

**Figure 1.**
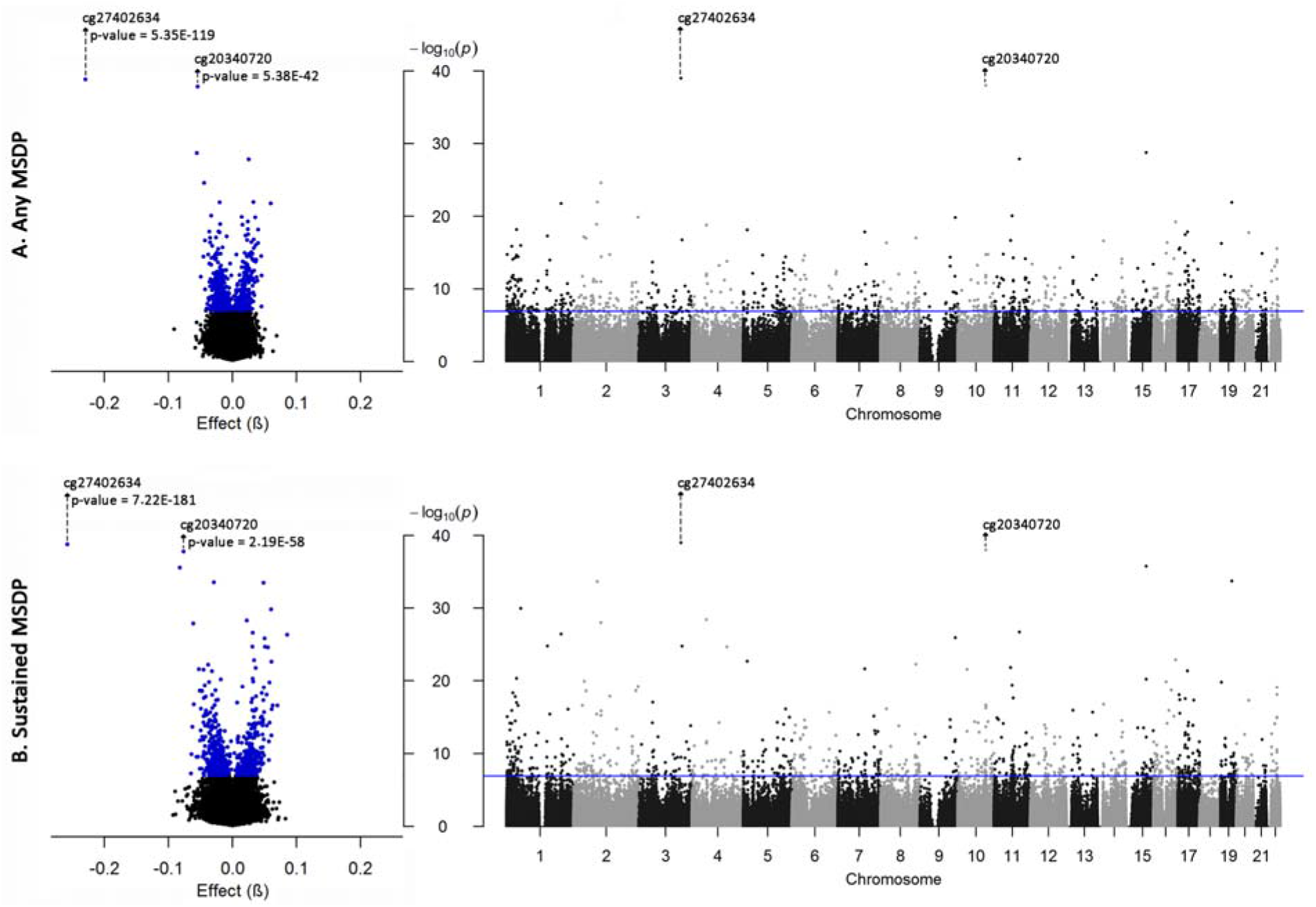
Volcano and Manhattan plots of the association between any MSDP (total N=1700, exposed =344) (A) and sustained MSDP (total N=795, exposed = 163) (B) with placental DNAm adjusted for maternal age, parity, maternal education and putative cellular heterogeneity. In the volcano plots, the x-axis shows the difference in DNAm (effect) with a possible range between 0 and 1, while the x-axis in the Manhattan plot represents genomic location; both plots share the same y-axis with –log_10_(*P*). Bonferroni thresholds for statistical significance are shown as blue dots and a blue horizontal line, for volcano and Manhattan plots, respectively. The y-axes were truncated to a minimum p-value of 1*10^-40^ (or maximum -log_10_(*P*) of 40), to allow for better visualization of the majority of our results. The CpGs that were impacted by y-axis truncation are indicated with arrows and annotated with their actual p-values.

The most notable association was observed at cg27402634, located upstream of the *LEKR1* gene and the non-coding RNA *LINC00886*, which showed the largest differential DNAm and smallest p-values in all meta-analyses. Placentas that were exposed to any MSDP had 22.95% lower DNAm (95% CI: 21.01-24.99% lower DNAm; p-value = 5.35E-119) and those exposed to sustained MSDP had 25.78% lower DNAm (95% CI: 24.02-27.54% lower DNAm; p-value = 7.22E-181) when compared to mothers that did not smoke at all during pregnancy. Though all cohorts observed substantial hypomethylation with MSDP at this CpG, the actual estimates of the associations were highly variable between cohorts for models of any MSDP (heterogeneity p-value = 2.66E-15), but relatively consistent for models of sustained MSDP (heterogeneity p-value = 1.55E-01) (Figure 2A). Overall, we consistency in the associations across cohorts for the vast majority of the 1224 CpGs that yielded Bonferroni-significant associations via meta-analysis: 92% and 97% of these CpGs yielded heterogeneity p-values > 0.01, for their associations with any and sustained MSDP respectively. In addition to cg27402634, we highlight those relationships that yielded that largest magnitudes of association: (|β_Any MSDP_|) > 0.05 for cg26843110 (*EDC3*), cg20340720 (*WBP1L*), and cg17823829 (*KDM5B*) (Figure 2B-2D). We identified numerous other noteworthy relationships but due to the large number of Bonferroni-significant associations from our meta-analyses, we highlight the relationship among the 20 most statistically significant CpGs from the primary meta-analysis of any MSDP going forward (Table 2).

**Figure 2.**
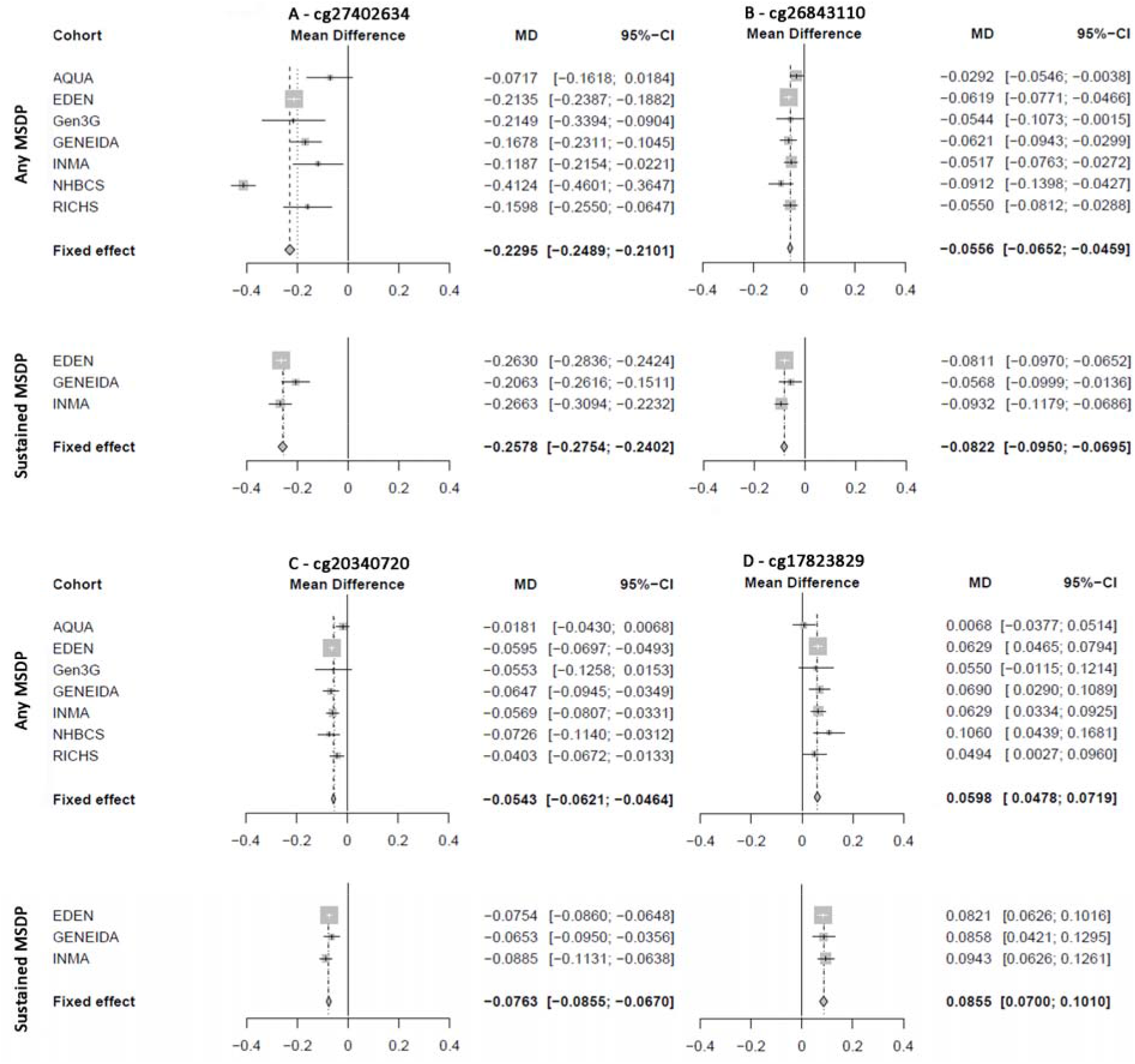
Forest plots of the cohort specific estimates of association and fixed-effect meta-analysis estimates of association between placental DNAm levels at (A) cg27402634, (B) cg26843110, (C) cg20340720, and (D) cg17823829 with any MSDP and sustained MSDP; models adjusted for maternal age, parity, maternal education, and putative cellular heterogeneity.

**Table 2.** Meta-analysis results from models of any and sustained MSDP, for the 20 CpGs with the smallest p-values for the model of any MSDP, and the percent increase in effect size between sustained and any MSDP (% Change); all models adjusted for maternal age, parity, education and putative cellular heterogeneity; CpGs that were not annotated with a gene name in the Illumina 450K annotation file have been annotated with their genomic region (ie. 4q12).

| CpG Annotations |  |  | Any MSDP |  |  | Sustained MSDP |  |  | % Change |
| --- | --- | --- | --- | --- | --- | --- | --- | --- | --- |
| CpG ID | Location | Annotated Gene (Region) | $\beta_1$ | S.E. | P-value | $\beta_1$ | S.E. | P-value | |
| cg17823829 | chr1:202765754 | *KDM5B* (Body) | 0.06 | 0.006 | 1.79E-22 | 0.086 | 0.008 | 3.80E-27 | 42.98 |
| cg26045080 | chr1:36807363 | *STK40* (Body) | 0.024 | 0.003 | 6.66E-19 | 0.032 | 0.003 | 4.86E-21 | 31.12 |
| cg00534380 | chr2:101766586 | *TBC1D8* (Body) | -0.044 | 0.004 | 2.53E-25 | -0.061 | 0.006 | 1.03E-28 | 38.46 |
| cg19246018 | chr2:240031588 | *HDAC4* (Body) | 0.015 | 0.002 | 1.39E-20 | 0.016 | 0.002 | 6.52E-20 | 8.28 |
| cg23752985 | chr2:85803571 | *VAMP8* (TSS1500) | -0.019 | 0.002 | 1.31E-19 | -0.025 | 0.003 | 3.77E-16 | 32.98 |
| cg00666842 | chr2:88366145 | *SMYD1* (TSS1500) | 0.033 | 0.003 | 1.17E-22 | 0.049 | 0.004 | 2.23E-34 | 48.17 |
| cg27402634 | chr3:156536860 | 3q25.31 | -0.23 | 0.01 | 5.40E-119 | -0.258 | 0.009 | 7.20E-181 | 12.33 |
| cg09491670 | chr4:53529646 | 4q12 | 0.016 | 0.002 | 1.60E-19 | 0.023 | 0.002 | 3.96E-29 | 43.31 |
| cg25585967 | chr5:14452105 | *TRIO* (Body) | 0.04 | 0.005 | 7.53E-19 | 0.061 | 0.006 | 2.16E-23 | 51.74 |
| cg12291408 | chr7:100037572 | 7q22.1 | -0.036 | 0.004 | 1.43E-18 | -0.052 | 0.005 | 2.34E-22 | 45.15 |
| cg14214914 | chr9:131870304 | *CRAT* (Body) | 0.036 | 0.004 | 1.57E-20 | 0.05 | 0.005 | 1.20E-26 | 41.29 |
| cg20340720 | chr10:104512523 | *WBP1L* (Body) | -0.054 | 0.004 | 5.38E-42 | -0.076 | 0.005 | 2.19E-58 | 40.52 |
| cg26648103 | chr11:66791718 | *SYT12* (5'UTR) | -0.033 | 0.004 | 8.98E-21 | -0.043 | 0.005 | 4.30E-20 | 30.51 |
| cg26115089 | chr11:93846406 | *HEPHL1* (3'UTR) | 0.025 | 0.002 | 1.33E-28 | 0.032 | 0.003 | 2.00E-27 | 24.8 |
| cg26843110 | chr15:74935742 | *EDC3* (Body) | -0.056 | 0.005 | 1.74E-29 | -0.082 | 0.007 | 1.74E-36 | 47.84 |
| cg26433445 | chr16:81764289 | 16q23.3 | 0.024 | 0.003 | 5.85E-20 | 0.034 | 0.003 | 1.34E-23 | 40.25 |
| cg24177452 | chr17:27494295 | *MYO18A* (5'UTR) | 0.025 | 0.003 | 3.40E-18 | 0.031 | 0.004 | 2.84E-18 | 27.35 |
| cg06716730 | chr17:35851459 | *DUSP14* (5'UTR) | -0.022 | 0.003 | 1.40E-18 | -0.032 | 0.003 | 4.47E-22 | 46.79 |
| cg03313447 | chr19:41829042 | *CCDC97* (3'UTR) | -0.02 | 0.002 | 1.26E-22 | -0.029 | 0.002 | 1.93E-34 | 45.73 |
| cg02341503 | chr20:45947123 | *ZMYND8* (Body) | -0.02 | 0.002 | 1.81E-18 | -0.025 | 0.003 | 4.83E-18 | 25.74 |

$\beta_1$ = Coefficient of the association between DNAm and MSDP; MSDP = Maternal smoking during pregnancy; S.E. = Standard Error.

### Expression Quantitative trait methylation (eQTM) analyses

We tested whether the DNAm levels at MSDP-associated CpGs were associated with the expression of nearby (± 250kb of the CpG) mRNA from 194 placental samples in the RICHS cohort, and corrected for multiple testing with the less conservative false discovery rate (FDR) due to the smaller sample size. We identified 170 associations between DNAm and gene expression within a 5% FDR (Excel Table S8), including 141 unique CpGs and 140 unique mRNAs; 72.4% of the eQTMs exhibited inverse associations, while inverse associations were most common and statistical significance was strongest for CpGs that were closer to the transcription start site (Supplemental Figure S3). Among the top 20 CpGs from our meta-analysis, eight were identified as eQTMs (5% FDR) (Table 3). Higher placental DNAm at cg27402634, was associated with lower expression (β = −2.32, and FDR < 5%) of *LEKR1*. Additionally, DNAm at cg17823829 (annotated to *KDM5B*) was most strongly associated lower expression of *PPFIA4* (FDR < 5%), and DNAm at cg26843110 (annotated to *EDC3*) was nominally associated with higher expression of *CSK*, while cg20340720 (annotated to *WBP1L*) was not associated with any nearby mRNA expression.

**Table 3.**
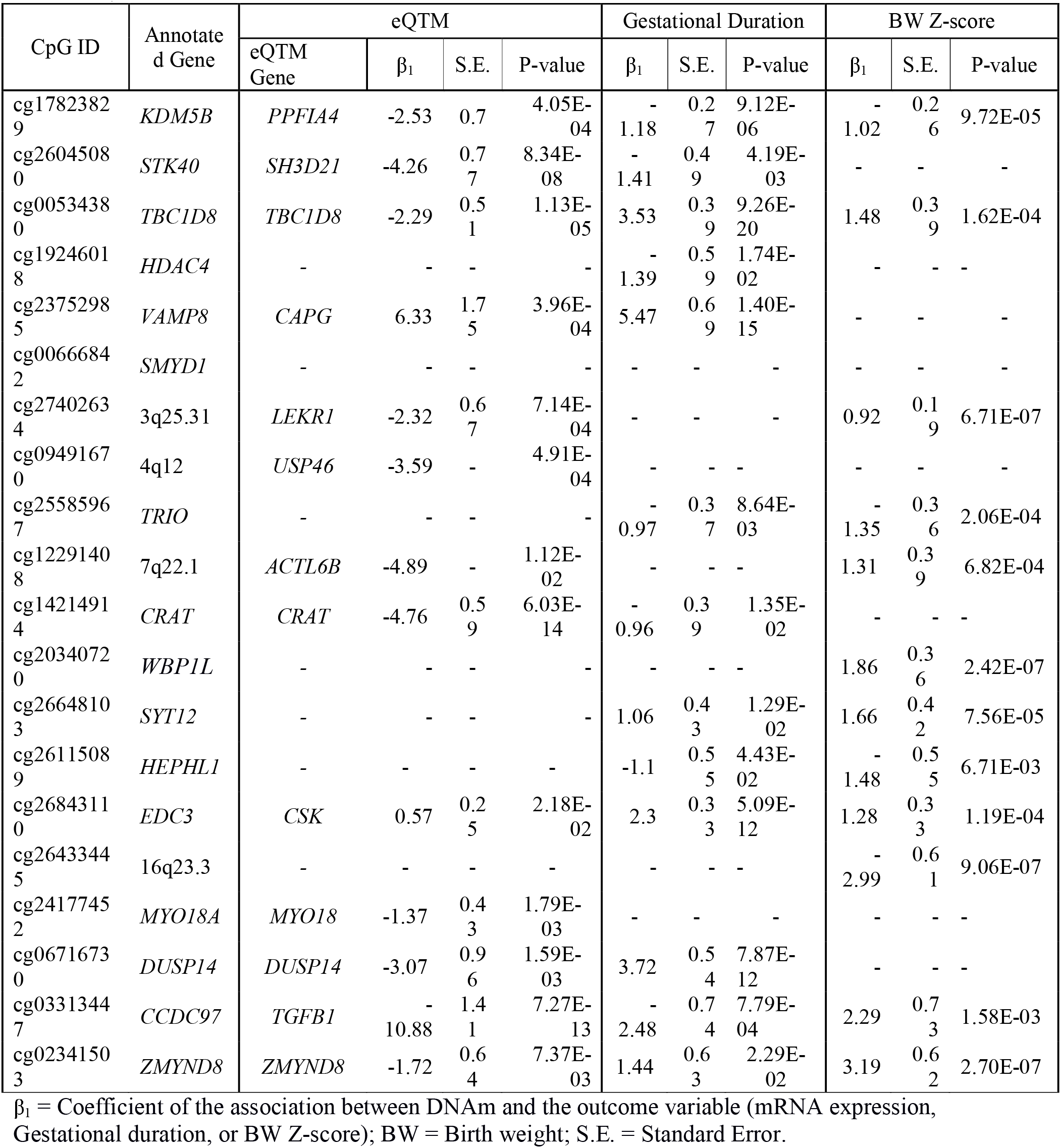
Results from eQTM models, DNAm versus gestational age at birth, and DNAm versus BW Z-score models that yielded at least nominally significant associations, among the 20 CpGs that yielded the most statistically significant associations with any MSDP in the primary meta-analysis (sorted in the same order as Table 2).

### Functional and regulatory enrichment analyses

Enrichment analyses were performed to gain insights into which biological processes may be impacted by these MSDP-associated CpGs. We performed gene-set enrichment analyses, which included 565 genes annotated to CpGs associated with any MSDP and 581 genes for sustained MSDP. 143 and 25 pathways were significantly (q-value < 0.05) enriched among CpGs associated with any MSDP (Excel Table S9) and sustained MSDP (Excel Table S10), respectively. In both gene-sets, “signaling by nerve growth factor (NGF)” was the most significant pathway (q-value = 2E-04 for any MSDP and q-value = 9.6E-03 for sustained MSDP). Other significantly enriched pathways were related to growth factors (VEGF, EGF, PDGF, IGF1R), hormones (aldosterone, insulin, TSH, GnRH), interleukins (IL2, IL4, IL7), myometrial contraction, vascular smooth muscle contraction, thrombin and platelet activation, signaling and aggregation. We also tested whether the genes annotated to MSDP-associated CpGs were enriched for regulatory regions of specific transcription factors (TFs). Most notably, our MSDP-associated CpGs were enriched for genes regulated by GATA1 and GATA2, as well as by RUNX1, TP63, SMAD4, AR, TP53 or PPARG (Excel Tables S11-S12).

We then examined whether the MSDP-associated CpG sites were enriched for CpG island locations, allele-specific germline differentially methylated regions (gDMR)^23^, regulatory features from the placenta specific 15-chromatin state annotation from ROADMAP^24^, or placenta specific partially methylated domains (PMD)^25^, which contain placenta-specific repressed genes (annotated to the results files in Supplementary Tables S3-S6). The MSDP-associated CpGs were depleted in PMDs (Supplemental Figure S4), highly enriched in placenta enhancers and depleted in transcription start sites and inactive states (Supplemental Figure S5). Additionally, MSDP-associated CpGs were depleted in CpG islands and shores, while enriched in CpG island shelves and open sea positions (Supplemental Figure S6). We identified 3 CpGs (cg05211790 and cg16360861 at *RAI14*, and cg05575921 at *AHRR* gene) that were within two candidate maternal gDMRs, but our overall set of MSDP-associated CpGs were neither depleted nor enriched for confirmed gDMRs (Supplemental Figure S7).

### Phenotype enrichment analyses

The genes annotated to our MSDP-associated CpGs were tested for phenotype enrichment using data from the database of Genotypes and Phenotypes (dbGAP). Our MSDP-associated genes were enriched for numerous phenotypes in dbGAP, including several adiposity phenotypes (body mass index (BMI), waist-hip ratio, and obesity), blood pressure, cardiovascular diseases, type 2 diabetes, asthma and respiratory function (Excel Tables S13-S14).

### Proximity to genetic variants for birth outcomes

We aimed to understand whether the MSDP-associated CpGs that we identified co-localize within the same genomic regions as genetic variants that have been associated with birth outcomes via genome-wide association studies (GWAS). Thus we investigated whether MSDP-associated CpGs were within ± 0.5 Mb (1 Mb window) of single nucleotide polymorphisms (SNPs) that have been associated with birth weight (BW), birth length (BL), head circumference (HC) and gestational age (GA)^26–31^, which have been added as annotations to the meta-analysis results files (Excel Tables S3-S6). Of the 324 birth outcome SNPs in autosomal chromosomes, 94 SNPs (83 loci) and 108 (97 loci) were within 0.5 Mb of CpGs that were associated with any or sustained MSDP, respectively (Excel Tables S15). Overall ∼16% of the 1224 MSDP-associated CpG sites were within 0.5 Mb of birth outcome SNPs, including cg27402634 (*LEKR1*), cg26843110 (*EDC3*) and cg20340720 (*WBP1L*). We also explored whether our MSDP-associated CpGs may be biased by methylation quantitative trait loci (mQTLs), in which SNPs influence the methylation levels at nearby CpGs. Two studies have examined this question in human placenta, identifying 866^32^ and 4,342^33^ placental mQTLs. Our findings did not appear to be biased by genetic variation as only 9 of the 1,224 MSDP-associated CpGs are previously characterized placental mQTLs.

### Association of DNAM at smoking associated loci and smoking related birth outcomes

We also performed a second meta-analysis to examine the relationships between DNAm with gestational age at birth, preterm birth, BW, BL, and HC z-scores, outcomes that are known to be related to maternal smoking. Of the 1224 CpGs tested, 341 (27.9%) were related to at least one birth outcome after Bonferroni-adjustment (0.05/1224). The majority of birth outcome associations were related to gestational age at birth (298 CpGs) (Excel Table S16). Preterm delivery, for which only two cohorts could contribute data (EDEN and NHBCS), produced similar associations found for GA, though fewer CpGs were statistically significant (Excel Table S17). We also found that numerous loci were associated birth size z-scores, with the majority of these being associated with BW (43 CpGs) (Excel Table S18), followed by BL (20 CpGs) (Excel Table S19) and HC (4 CpGs) (Excel Table S20). Some of the CpGs associated with GA were also associated with birth size measurements, even though they were standardized for GA, suggesting independent associations with both gestational duration and fetal growth (Supplemental Figure S8A), including 6 CpGs (annotated to *SYNJ2*, *PXN*, *PTPRE*, *IGF2BP2*, 4q21.1) shared between GA and BW, and 2 (annotated to *POLR3E* and *LOC441869*) shared between GA and BL. Among the CpGs that yielded at least one Bonferroni-significant association with birth outcomes, CpGs that tended to have positive associations with birth outcomes clustered together and were typically hypomethylated with MSDP, while CpGs that exhibited inverse associations with birth outcomes tended to be hypermethylated with exposure to MSDP (Supplemental Figure S8B).

Among our top 20 CpGs that were associated with any MSDP, 5 were associated with GA at birth and 4 were associated with BW z-scores after Bonferroni adjustment (Table 3). DNAm at cg27402634 (*LEKR1*) and cg20340720 (*WBP1L),* both located close to BW-SNPs and for which MSDP associated with lower DNAm, were associated with larger BW (p-value = 6.71E-07, and p-value = 2.42E-07). On the other hand, DNAm at cg26843110 (*EDC3;* hypomethylated in response to MDSP and also close to BW-SNPs) and at cg17823829 (*KDM5B;* hypermethylated) were associated with longer and shorter gestational ages at birth, respectively (p-value = 5.09E-12, and p-value = 9.12E-06). Forest plots of BW z-scores and GA for these four CpGs are shown in Figure 3.

**Figure 3.**
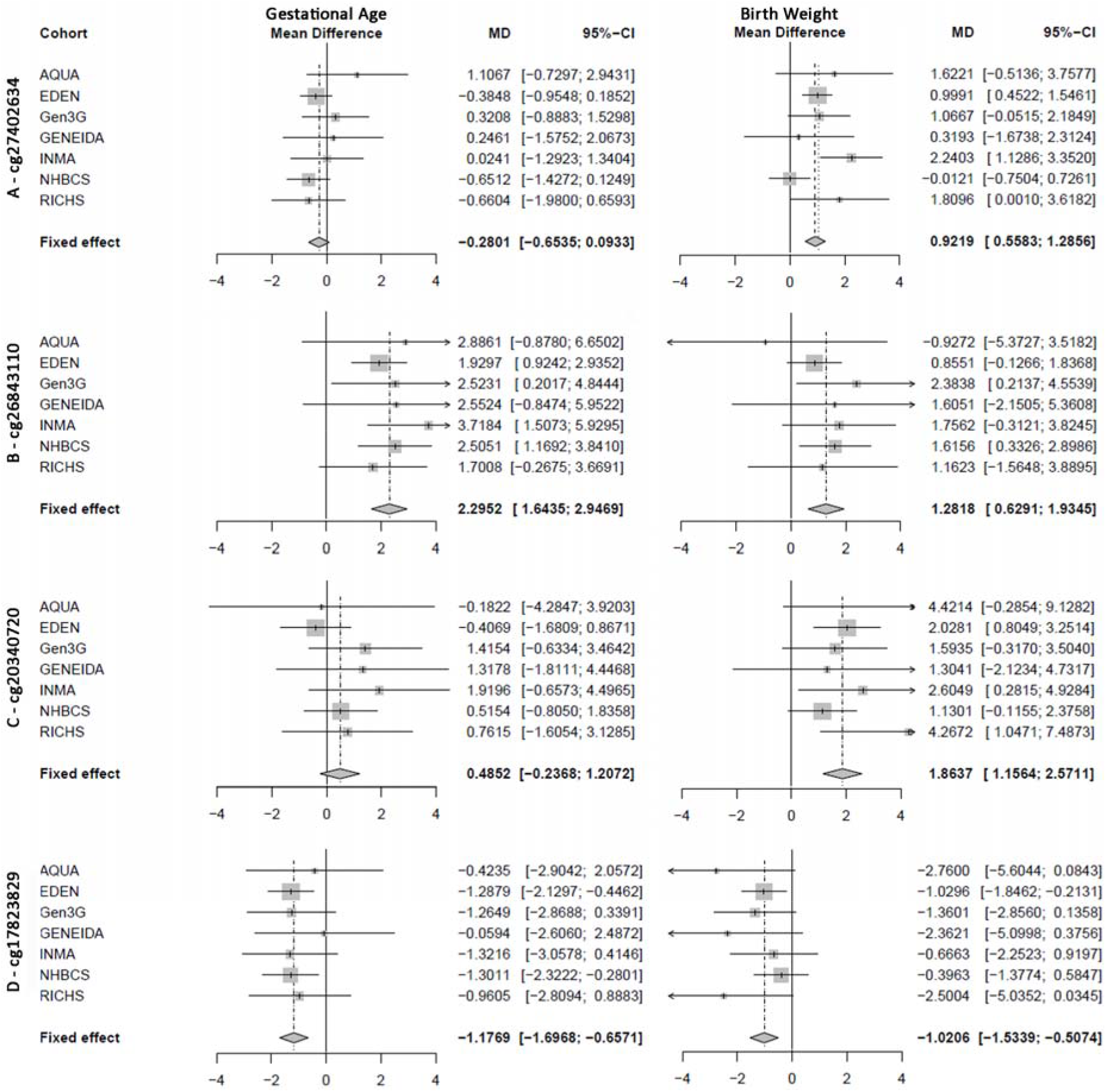
Forest plots of the cohort specific estimates of association and fixed-effect meta-analysis estimates of association between higher levels of placental DNAm at (A) cg27402634, (B) cg26843110, (C) cg20340720, and (D) cg17823829 with gestational age and birth weight z-scores; models adjusted for maternal age, parity, maternal education, and putative cellular heterogeneity.

We summarize the results for all secondary analyses with a circos plot, for those 548 CpGs that yielded Bonferroni significant associations with both any and sustained MSDP (Figure 4). Among these, 21 CpGs were associated both with mRNA expression (FDR < 5%) and at least one birth outcome (Bonferroni-significant), which have been annotated with the gene symbols from their respective eQTM models.

**Figure 4.**
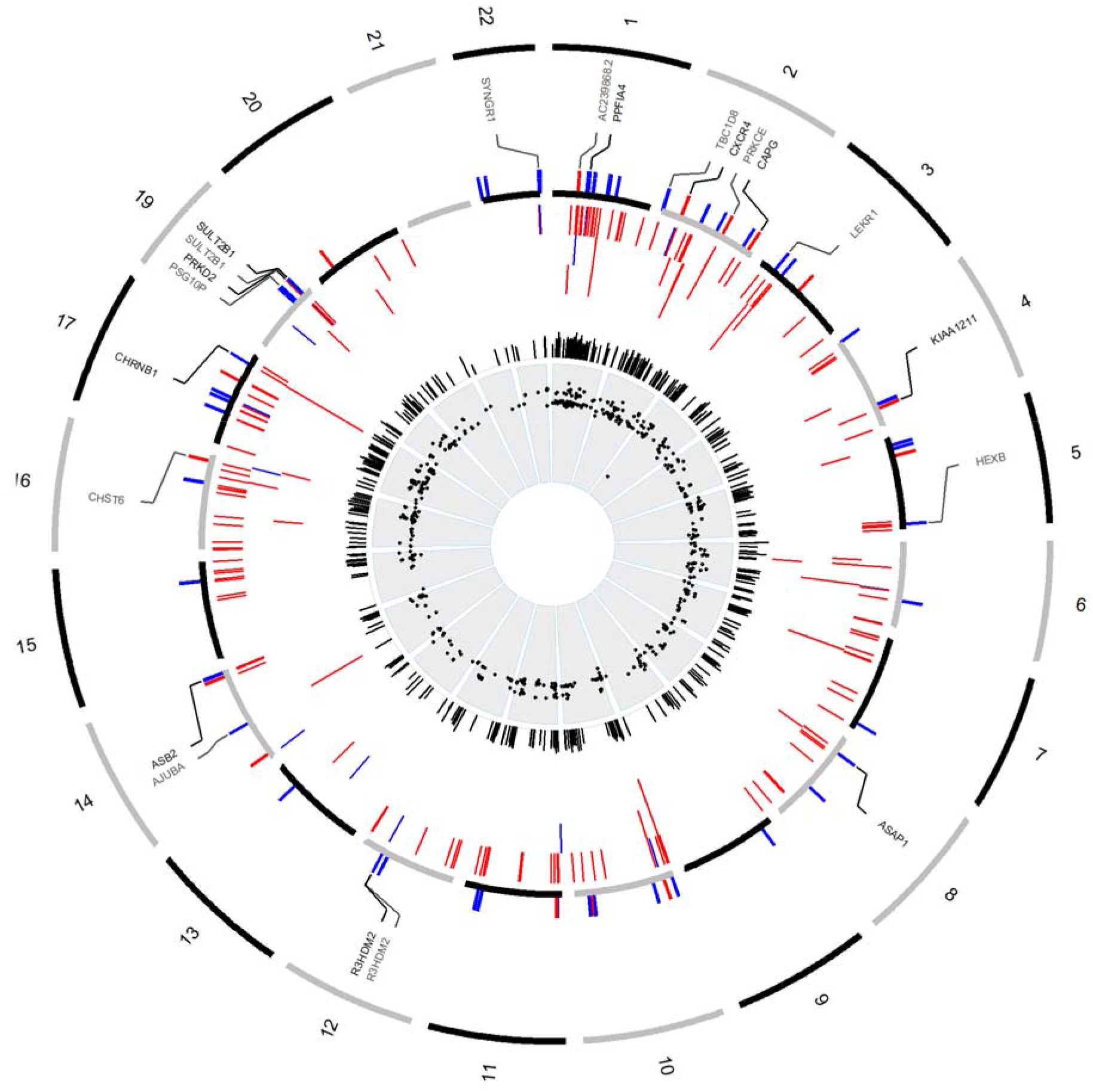
Circos plot summarizing all analyses for the 548 CpG sites that yielded Bonferroni-significant associations with both any and sustained MSDP. From inner to outer-band: Black points = differential DNAm between any- and no MSDP; Black Bars = percent increase in magnitude of association when comparing results from any MSDP to sustained MSDP; Color Bars = Positive (red) or Inverse (blue) associations between DNAm and HC, BL, BW, and GA, that yielded Bonferroni-significant associations; Color Bars = Positive (red) or Inverse (blue) associations between DNAm and nearby gene expression, that yielded an association at a 5% FDR; Names of eQTM genes were annotated to the outer band for CpGs that yielded an association between DNAm and expression with a 5% FDR and yielded an association between DNAm and at least one birth outcome after Bonferroni-adjustment.

### Comparison with smoking-sensitive CpGs in cord blood

We assessed whether the DNAm signatures of MSDP in the placenta were consistent with MSDP associations in cord blood previously reported by the PACE consortium^10^. Only nine CpGs annotated to seven unique genes (*AHRR, CYP1A1*, *GNG12*, *PXN*, *RNF122*, *SLC23A2*, and *ZBTB4*) yielded Bonferroni-significant associations in both tissues, out of 1224 CpGs from our study and 568 CpGs from the cord blood study (Table 4). Of note, the CpGs within *CYP1A1* and *RNF122* showed opposite directions of association with MSDP in cord blood and placenta. We also compared the parameter estimates from our study that yielded associations with MSDP within 5% FDR to the parameter estimates of those CpGs described in cord blood also within a 5% FDR^10^. There was no overall correlation (r^2^<0.1) of the regression coefficients across these two tissues (Supplemental Figure S9).

**Table 4:**
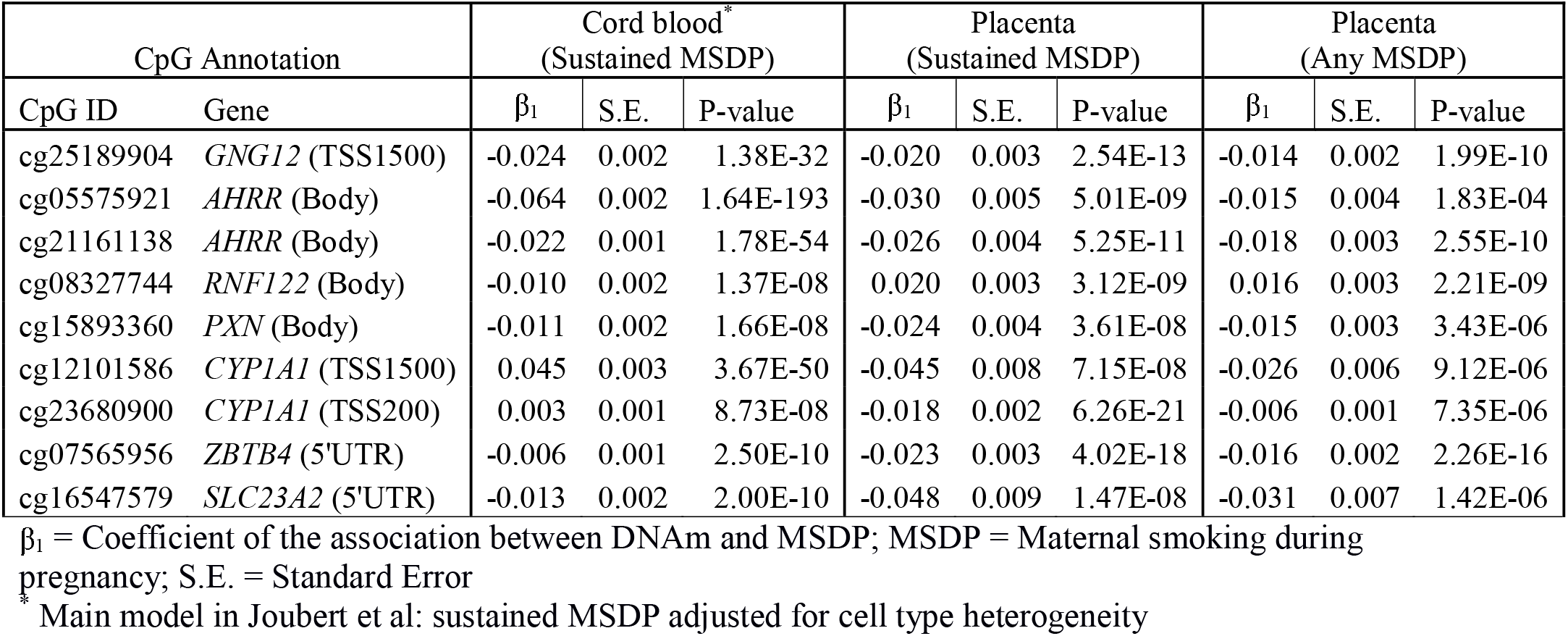
Differentially methylated CpGs with MSDP in placenta (from our meta-analysis) and cord blood (from a published PACE meta-analysis^10^) that yielded Bonferroni-significant associations in both tissues.

## Discussion

We identified 1224 CpG sites with placental methylation levels that were associated with any or sustained MSDP. Differential DNAm was greater with increased duration of exposure at all loci that were associated with both any and sustained MSDP, and a large proportion of the MSDP-associated CpGs were related to birth outcomes. Those CpGs that were observed to have higher DNAm associated with MSDP, tended to be inversely associated with gestational age and birth size, while CpGs exhibiting lower DNAm with MSDP tended to be positively associated with gestational age and birth size.

The MSDP-associated loci with the most statistically significant association (cg27402634), also identified in prior smoking EWAS of placental tissues^15^, yielded dramatically lower DNAm levels in association with MSDP exposure. This effect-size is much larger in magnitude (∼25% difference for sustained MSDP) compared to what has generally been observed in most exposure-focused EWAS, though within the same range as a CpG site in *AHRR* (cg05575921; 18% difference between exposed and unexposed) from a prior EWAS of current smoking and blood DNAm^34^. Additionally, decreased placental DNAm at cg27402634 correlates with increased expression of *LEKR1,* and associates with smaller BW and BL. Thus MSDP-associated hypomethylation at this CpG would be consistent with the well-known effect of maternal smoking resulting in shorter gestation and smaller birth size.

The functional activities of cg27402634, or corresponding *LEKR1* gene, in human placental tissues are not known. However, GWAS findings provide evidence that genetic variants within this region might be involved in fetal growth and possibly metabolic programming. For instance, the SNP rs1482852 or its proxies (rs900400; rs13322435) have been associated with different parameters of fetal growth^35^, adiposity in newborns^36,37^, maternal adiponectin levels, cord blood leptin^37^, and insulin release after an oral glucose challenge^38^. These findings from genetic studies in combination with our current study, implicate that this locus on chromosome 3 (3q25.31) contains very active determinants of growth regulation and metabolic activity, and that placental DNAm at cg27402634 is highly responsive to maternal smoking. Future mechanistic work is necessary to investigate whether the placental epigenetic regulation at this locus specifically influences placental functions and/or overall growth and metabolic functions of the developing fetus.

We identified numerous other notable MSDP-associated loci in addition to those of cg27402634 (*LEKR1*), and highlight those CpGs yielding the strongest magnitudes of effect (cg20340720, cg26843110, and cg17823829). MSDP was associated with lower DNAm at cg20340720, located within *WBP1L* (also annotated as *C10orf26*), while lower DNAm at this CpG correlated with lower with BW and BL z-scores. Genetic variants nearby to this CpG have been related to BW^31^ and blood pressure^39^. We also observed lower DNAm with MSDP at cg26843110, which is within the body of the *EDC3* gene, and is nearby to SNPs associated with BW (rs3784789^31^). Lower DNAm at cg26843110 associated with shorter GA at birth, and decreased expression of *CSK*, which is involved in trophoblast differentiation^40^ as well as blood pressure and aldosterone regulation^41^. Finally, cg17823829 (annotated to *KDM5B*) was hypermethylated with MSDP. Higher DNAm at this CpG correlated with shorter GA at birth and with lower expression of *PPFIA4* gene, which can be induced in response to hypoxia^42^.

Our enrichment analyses identified numerous pathways that are critical to placental growth and development, such as vascularization, hormone signaling and inflammatory cytokines. Multiple pathways involving vascular endothelial growth factors (VEGF) and nerve growth factors (NGF) populated the top of our enrichment lists. The VEGFs and their receptors are required for all steps of placental vascularization^43^, while nerve growth factor (NGF) modulates immune activity, inflammation and angiogenesis in the placenta^44^. Thus, our findings may be related to perturbed placental vascularization or angiogenic signaling. Altered placental vasculature is the most common placental pathology identified in pregnancy complications^43^, and MSDP can result in placental vascular remodeling^5^. Our findings may represent, in part, an epigenetic footprint of the placental vascular and angiogenic response to MSDP. Our CpGs were also enriched for genes regulated by specific transcription factors, most notably GATA1 and GATA2. Together with PPARG and TP63, GATA factors are part of the core transcriptional regulatory circuit that guides and maintains proper trophoblast differentiation^45,46^. Placentas lacking *PPARG* have lethal defects in placental vascularisation^43^, and angiogenic activity is reduced in placentas lacking *GATA2*^47^. Additionally, our MSDP-associated CpGs were enriched in placental enhancers, while depleted in transcription start sites, inactive states, PMDs^25^ and gDMRs^23^, overall suggesting that the CpG sites that we identified are located in active regulatory regions.

Genes that are annotated to the CpGs that we studied have been linked to human health and disease traits via dbGAP, including a number of conditions that are part of the metabolic syndrome (BMI, obesity, cholesterol, blood pressure), for which prior links to MSDP have been reported^48,49^. This may indicate that the placental genes whose regulation is impacted by MSDP, are involved in energy uptake and expenditure, lipid and glucose metabolism, blood pressure regulation, and inflammation, which are some of the key physiological processes that are disrupted in the pathogenesis of metabolic syndrome^50^. The MSDP-associated CpGs were also enriched for genes linked to asthma and impaired respiratory function, which are known to be caused by MSDP^51^, but it is unclear if altered regulation of these genes in the placenta has consequences on the development of the respiratory system. Furthermore, 128 of the 324 SNPs that have previously been associated with birth size or gestational age at birth^26–31^ were in similar genomic proximity (within 0.5 Mb) to our CpGs, suggesting the MSDP-associated differential methylation in the placenta occurs in regions of the genome that are heavily involved in growth and development. Many of these SNPs have been shown to be related to birth outcomes, glycemic traits, blood pressure, and height, while the largest GWAS of BW to date concludes that the link between lower BW and later cardio-metabolic traits are largely driven by shared genetic effects^31^. Future research is needed to characterize this convergence of genetic effects on growth and placental epigenetic responsiveness to MSDP within similar genomic proximities. It is possible that MSDP, genetic variation, and placental DNAm in these regions yield additive or interactive effects on birth outcomes; prior studies have addressed this question in blood^52,53^. Additionally, while our MSDP-associated CpGs are in similar genomic regions (1 Mb windows) with birth outcome SNPs, very few of our CpGs (9 of 1224) and very few of the birth outcome SNPs (3 of 324) are known placental mQTLs^32,33^.

We compared our findings to those of a previous PACE meta-analysis of MSDP and cord blood DNAm^10^. Two of the CpGs that we identified within the *AHRR* gene (cg05575921 an eQTM for *AHRR* in placenta, and cg21161138), have been consistently observed to be sensors of MSDP exposure in cord blood. Only four other CpG sites, (annotated to *GNG12*, *PXN*, *ZBTB4*, and *SLC23A2*) were differentially methylated in both placenta and cord blood with the same direction of association in both tissues. Additionally, three CpG sites within *CYP1A1* and *RNF122* were identified in both meta-analyses but with different directions of association in cord blood versus placenta. Interestingly, we observed *CYP1A1* to be hypomethylated in placenta with exposure to MSDP, which is consistent with studies of adipose, skin, and lung tissues^54^, but this CpG was hypermethylated in the cord blood meta-analysis^10^. Additionally, the most statistically significant association with MSDP in placenta (cg27402634, *LEKR1*) was not associated with MSDP in the cord blood meta-analysis. These observations, and the lack of overall correlation in regression coefficients when comparing placental and cord blood responses to MSDP, suggest that there are unique tissue-specific molecular responses to this exposure.

The above findings should be interpreted within the context this study’s limitations. MSDP was self-reported and subject to misclassification, though differential misclassification likely would have biased our findings towards the null. We modeled two different definitions for MSDP and found that the models with sustained MSDP produced larger magnitudes of association, as has been previously found in studies of blood DNAm^55^. Although these findings suggest that increased duration of MSDP is associated with greater differential DNAm, we did not assess dose-response patterns (ie. number of cigarettes or cotinine concentrations), which should be the focus of future investigations. Our study predominantly consisted of samples from mother-infant pairs of European ancestry, and thus additional studies involving diverse racial and ethnic backgrounds are needed in order to improve the generalizability of these findings. While it is unlikely that placental DNAm would influence maternal smoking, the observed associations between DNAm levels and reproductive outcomes could be due to reverse causation. Placenta is a heterogeneous tissue with multiple different cell types^56^ that serve different functions and thus have different epigenetic states^57^. To correct for this, we estimated and adjusted for variability in placental DNAm that may be due to tissue heterogeneity. We utilized a data driven approach that was not based on a methylome reference, as no references for placental cell-type methylomes are currently available. Adjustments for these estimates of putative cellular admixtures did reduce heterogeneity in the meta-analyses, thus improving the consistency of observed associations between MSDP and DNAm across these independent cohorts. However, it is possible that residual confounding may have influenced some of our results.

Despite these limitations, our study had numerous strengths, including a large overall sample-size and seven independent studies to identify these relationships. We used harmonized definitions of exposure variables and covariates, standardized protocols for quality control and pre-processing of DNAm data, and standardized methods for estimating/adjusting for tissue heterogeneity and for statistical analyses. We also performed secondary analyses involving mRNA expression, functional and phenotype enrichment, overlap with GWAS hits for reproductive outcomes, and meta-analyses of DNAm variation with birth outcomes to provide biological and health-related interpretations of our findings.

We identified a DNAm signature of MSDP in the placenta that shows substantial differences from that observed in cord blood, most notably the CpG in close proximity to *LEKR1*. Many of the identified MSDP-associated loci are involved in biological process that are known to play critical roles in placental development, including vascularization, angiogenesis, and inflammation. Additionally, a large proportion of the MSDP-associated CpGs were also associated with GA at birth, or birth size z-scores, suggesting these placental epigenetic variations may be intermediate molecular markers between MSDP and these outcomes. Further study is required to determine whether these epigenetic variations are causal mediators within these relationships or reflecting other processes.

## Material and methods

### Participating cohorts

Cohorts that are members of the PACE consortium were identified for participation in the current study if they had existing DNAm data quantified from placental tissue via the Illumina Infinium HumanMethylation450 BeadChip and if they had obtained information on self-reported smoking during pregnancy. The seven cohorts that contributed to the meta-analysis of any MSDP included AQUA^16^, EDEN^17^, Gen3G^18^, GENEIDA, INMA^19^, NHBCS^20^ and RICHS^21^. EDEN, GENEIDA and INMA also contributed to the sustained MSDP stratified analyses. RICHS contributed RNAseq data for analyses with mRNA expression. All cohorts acquired ethics approval and informed consent from participants prior to data collection through local ethics committees. Exclusion criteria for this study were: non-singleton births, pre-eclampsia and DNAm not assessed in the fetal side of the placenta. All participants in the study were of European ancestry, except 1.85% of EDEN mothers. Detailed methods for each cohort are provided in the Supplementary Material (Supplemental Methods File).

### Tobacco smoking definitions

Any MSDP was defined as “yes” if mothers reported smoking cigarettes at any time during pregnancy. Sustained MSDP was defined as “yes” when mothers reported smoking cigarettes at least in the 1^st^ and 3^rd^ trimester of pregnancy. For both exposure variables, the comparison group was defined as the mothers that reported no smoking during any of the pregnancy.

### Placental genome-wide DNAm data acquisition, quality control and normalization

Placental DNAm from the fetal side was assessed with the Infinium Human-Methylation450 array (Illumina, San Diego, CA USA). See Supplementary Methods file for extra details on placenta collection, DNA extraction and DNAm acquisition in each cohort. Quality control of DNAm was standardized across all cohorts. Low quality samples were filtered out and probes with detection p-values > 0.01 were excluded. Beta-values were normalized via functional normalization^58^ and beta-mixture quantile normalization (BMIQ)^59^ was applied to correct for the probe type bias. Cohorts applied ComBat to remove batch effects when applicable. Probes that hybridize to the X/Y chromosomes, cross-hybridizing probes and probes with SNPs at the CpG site, extension site, or within 10 bp of the extension site with an average minor allele frequency > 0.01 were filtered out^60^. Overall, 418,639 probes and 415,396 were available for modelling any MSDP and sustained MSDP, respectively. Finally, DNAm extreme outliers (<25^th^ percentile – 3*IQR or >75^th^ percentile + 3*IQR across all the samples) were trimmed.

### Estimates of putative cellular heterogeneity

Placental putative cellular heterogeneity was estimated from DNAm data using a reference-free cell-mixture decomposition method^61^. The number of surrogate variables ranged from 2 to 5 depending on the cohort. Models for differential DNAm were corrected for the number of surrogate variables minus one to reduce multi-collinearity.

### Genome-wide differential DNAM analyses

Within each cohort, robust linear regression from the “MASS” package^62^ in R were used to account for potential heteroskedasticity while testing the associations between normalized DNAm beta values at each CpG with any MSDP and sustained MSDP. Models were adjusted for maternal age, parity, maternal education and cohort-specific variables first unadjusted for putative cellular heterogeneity then adjusted for cellular heterogeneity. We performed inverse variance-weighted fixed-effects meta-analyses using METAL^63^. The meta-analysis was performed independently by two groups to ensure consistent results. CpGs not retained in at least 2 cohorts were filtered out. We used the Bonferroni adjustment to control for multiple testing. To examine whether increased duration of exposure (sustained smoking versus any smoking) yielded increased magnitudes of association, we calculated the percent change in the coefficients between the two models (|β_Sustained_| - |β_Any_|)/|β_Any_| * 100. Secondary analyses were only performed on CpGs that yielded Bonferroni significant associations with any or sustained MSDP in models that were adjusted for putative cellular heterogeneity.

### Expression quantitative trait methylation (eQTM) loci

We performed expression quantitative trait methylation (eQTM)^64^ analyses in the RICHS cohort. Transcription was measured via RNA-seq on 194 placentas. The details of sample collection, assay, and QC for the RNA-seq data are presented in detail elsewhere^65^, and summarized in the Supplementary Material (Supplementary Methods File). In this dataset, we identified 6523 unique transcripts annotated to an Ensembl ID (GrCh37/hg19) and with a transcriptional start site (TSS) within 250 kb upstream or downstream of 1184 out of the 1224 candidate CpGs. The association between DNAm and expression levels was assessed via 10295 linear regression models using the MEAL package^66^ in R. We report the results for all models yielding nominally significant associations (raw p-values <0.05), statistically significant eQTMs were determined at 5% FDR.

### CpG site annotation

We annotated CpGs to genes and CpG islands with notations from the Illumina HumanMethylation 450K annotation file, and with several regulatory features using publicly available data: placental 15-chromatin states^67^ released from the ROADMAP Epigenomics Mapping Consortium^24^ (ChromHMM v1.10), placental partially methylated domains (PMDs)^25^ and placental germline differentially methylated regions (gDRMs)^23^.

### Enrichment analyses

Functional enrichment analyses were performed at the gene level via ConsensusPathDB^68^ using KEGG, Reactome, Wikipathways, Biocarta as reference gene-sets. ConsensusPathDB performs a hypergeometric test and corrects multiple-testing with FDR. Enrichment for transcription factors and for phenotypes were assessed at the gene level with EnrichR using ENCODE and ChEA consensus TFs from ChIP-X database, and dbGaP database, respectively. EnrichR results were ranked using the combined score (P-value computed using Fisher exact test combined with the z-score of the deviation from the expected rank).^69^ Enrichment for regulatory features was assessed with the hypergeometric test, and P-values were Bonferroni-corrected for 15 (placental chromatin 15-states) and 6 (relation to CpG island) tests, respectively.

### Overlap of MSDP-sensitive CpG sites and birth outcome SNPs

Co-localization between MSDP-associated CpGs in placenta with previously identified BW, BL, HC and GA SNPs from the largest genome-wide association studies (GWAS) to date^26–31^ was assessed using the GenomicRanges package in R^70^. We identified which CpGs were located within 1 Mb windows (± 0.5 Mb) surrounding each of the 324 autosomal SNPs, which correspond to 280 potential unique loci. Unique loci were defined based on the criteria in Warrington et al. 2019^31^, and linkage disequilibrium in Europeans (r^2^ > 0.1 in < 2Mb).

### Association between DNAm and birth outcomes

Within each cohort, robust linear regression models were utilized to test the association between normalized DNAm beta values at each CpG as the independent variable and gestational age at birth (inverse normal transformation of sex residuals), BW z-scores, BL z-scores, and HC z-scores as the dependent variables. Logistic regression was used to examine the relationships between DNAm and pre-term birth (defined as <37 weeks of gestation). Birth size z-scores were calculated using international references from the INTERGROWTH-21^st^ Project^71^ and standardized by both gestational age and newborn sex. Models were adjusted for maternal age, parity, maternal education, cohort-specific variables (see Supplemental Methods) and putative cellular heterogeneity. Inverse variance-weighted fixed-effects meta-analyses^63^ were again used to estimate pooled associations. Multiple testing was controlled with the Bonferroni adjustment (α = 0.05/1224).

### Comparison of MSDP-sensitive CpGs in placenta with cord blood

We examined the consistency between MSDP-sensitive CpGs in placenta and in cord blood^10^. First we compared the coefficients from the models for sustained MSDP in cord blood, unadjusted for cellular heterogeneity, to results for both any and sustained MSDP in placenta, adjusted for cellular heterogeneity, using Pearson correlation coefficients.

All DNAm data processing and analyses were conducted in R, with the exception of the meta-analyses which were performed with METAL.

## Supporting information

Supplemental Contents and Methods

## Acknowledgements

We would like to thank all the families that participated in these studies for their generous contribution. Detailed acknowledgements and funding can be found in Supplementary Material.

## Author contributions

TME, MVU, CJM, JL, MFH and MB conceived of and designed the study. Study-specific analyses were completed by JMC, (AQUA), ES (EDEN), AC (Gen3G), PCS and JMM (GENEIDA), MVU (INMA) and TME (NHBCS and RICHS). MV meta-analysed these results. TME, MV and MB performed the follow-up analyses. TME, MVU, JS, CJM, MFH, and MB interpreted the results. TME, MVU, and MB wrote the first draft of the manuscript. All authors (TME, MVU, ES, AC, ML, JMC, CL, ERB, NFJ, BH, PP, BGA, JH, MAD, MRK, CI, LB, PCS, YJL, KH, TB, MAC, JMM, EM, JC, MFF, JT, AGM, SJL, JS, CJM, JL, MFH, MB) read and critically revised drafts of the manuscript. Correspondence can be addressed to TME and MB.

