## Supplemental Contents and Methods for "Placental DNA methylation signatures of maternal smoking during pregnancy and potential impacts on fetal growth"

**Excel Table S1.** Distribution of maternal smoking during pregnancy, demographic variables, birth outcomes, and covariates, by cohort.

**Excel Table S2.** Proportion of maternal smokers that were included in each model, GIF and number of significant CpGs for each model, for the meta-analyses and for individual cohorts; GIF - Genomic Inflation Factor.

**Excel Table S3.** Meta-analysis results for the association between any MSDP and placental DNAm, while controlling for maternal age, parity, and maternal education, but not adjusted for putative cellular heterogeneity; included data from AQUA, EDEN, Gen3G, GENEIDA, INMA, NHBCS, & RICHS.

**Excel Table S4.** Meta-analysis results for the association between sustained MSDP and placental DNAm, while controlling for maternal age, parity, and maternal education, but not adjusted for putative cellular heterogeneity; included data from EDEN, GENEIDA & INMA.

**Excel Table S5.** Meta-analysis results for the association between any MSDP and placental DNAm, while controlling for maternal age, parity, maternal education, and putative cellular heterogeneity; included data from AQUA, EDEN, Gen3G, GENEIDA, INMA, NHBCS, & RICHS.

**Excel Table S6.** Meta-analysis results for the association between sustained MSDP and placental DNAm, while controlling for maternal age, parity, maternal education, and putative cellular heterogeneity; included data from EDEN, GENEIDA & INMA.

**Excel Table S7.** Results from any MSDP meta-analysis when only EDEN, GENEIDA, and INMA were included for the 548 loci that were Bonferroni-significant for associations with both any and sustained MSDP; difference in estimated differential methylation between models for any and sustained MSDP (from original analysis with all cohorts, and secondary model only including EDEN, GENEIDA, and INMA).

**Excel Table S8.** Association between methylation levels at Bonferroni significant CpG sites (associated with any and/or sustained MSDP) vs. expression levels of mRNA within ± 250 kb of the CpG site.

**Excel Table S9.** Enrichment for functional pathways among CpGs associated with any MSDP, annotated to closest gene from the Illumina annotation file.

**Excel Table TS10.** Enrichment for functional pathways among CpGs associated with sustained MSDP, annotated to closest gene from the Illumina annotation file.

**Excel Table TS11.** Enrichment for transcription factor motifs for CpGs associated with any MSDP, annotated to closest gene from the Illumina annotation file.

**Excel Table TS12.** Enrichment for transcription factor motifs for CpGs associated with sustained MSDP, annotated to closest gene from the Illumina annotation file.

**Excel Table TS13.** Enrichment for phenotypes among CpGs associated with any MSDP, annotated to closest gene from the Illumina annotation file.

**Excel Table TS14.** Enrichment for phenotypes among CpGs associated with sustained MSDP, annotated to closest gene from the Illumina annotation file.

**Excel Table TS15.** Maternal and fetal SNPs associated with birth size and gestational age and their proximities to CpGs associated with any and sustained MSDP in placenta and with sustained MDSP in cord blood.

**Excel Table TS16.** Meta-analysis results for the association between placental DNAm and gestational age (inverse normal transformation of sex residuals, in days) adjusted for maternal age, parity, and maternal education, and putative cellular heterogeneity; included data from AQUA, EDEN, Gen3G, GENEIDA, INMA, NHBCS & RICHS.

**Excel Table TS17.** Meta-analysis results for the association between placental DNAm and preterm delivery (< 37 weeks gestation) adjusted for maternal age, parity, and maternal education, and putative cellular heterogeneity; included data from EDEN & NHBCS.

**Excel Table TS18.** Meta-analysis results for the association between placental DNAm and birth weight z-score (standardized by sex and gestational age via international reference), adjusted for maternal age, parity, and maternal education, and putative cellular heterogeneity; included data from AQUA, EDEN, Gen3G, GENEIDA, INMA, NHBCS & RICHS.

**Excel Table TS19.** Meta-analysis results for the association between placental DNAm and birth length z-score (standardized by sex and gestational age), adjusted for maternal age, parity, and maternal education, and putative cellular heterogeneity; included data from EDEN, Gen3G, GENEIDA, INMA, NHBCS & RICHS.

**Excel Table TS20.** Meta-analysis results for the association between placental DNAm and head circumference z-score (standardized by sex and gestational age), adjusted for maternal age, parity, and maternal education, and putative cellular heterogeneity; included data from EDEN, Gen3G, GENEIDA, INMA, NHBCS & RICHS.

**Figure S1.** QQ-plot for the meta-analyses of the association between placental DNAm and any MSDP without (A) and with adjustment for cellular heterogeneity (B) and sustained MSDP without (C) and with adjustment for cellular heterogeneity (D); N=1700 (A and B) and N=795 (C and D).

**Figure S2.** Plots showing the increase in the magnitude of coefficients for models of sustained MSDP versus any MSDP, (A) when the meta-analysis included all participating cohorts, and (B) when meta-analysis only included those cohorts that also participated in models of sustained MSDP (EDEN, GENEIDA and INMA); this analysis included the 548 CpGs that were Bonferroni-significant for associations with any and sustained MSDP.

**Figure S3**. –log10**(**P-value) versus distance between gene transcription start site (TSS) and CpG site for 1219 eQTMs. In blue positively associated eQTMs at 5% FDR, and in red inversely associated eQTMs at 5% FDR.

**Figure S4.** Proportion of CpGs annotated to placental partially methylated domains (PMDs) for those CpGs associated with MSDP.

**Figure S5.** Proportion of CpGs annotated to each of the 15 placenta chromatin states from ROADMAP, by significance and direction of the effect for associations with any MSDP (A) and sustained MSDP (B).

**Figure S6.** Proportion of CpGs annotated to each of the 6 locations in relation to a CpG island, by significance and direction of the effect, for those CpGs associated with any (A) and sustained (B) MSDP.

**Figure S7.** Proportion of CpGs annotated to candidate (A) or confirmed (B) placental germline differentially methylated regions (gDMR) that were associated with MSDP.

**Figure S8.** Venn diagram showing overlaps in CpGs that were associated with gestational age (GA), birth weight (BW), birth length (BL) and head circumference (HC) at Bonferroni significance (1224 CpGs tested; 339 significantly associated with birth outcomes), and (B) Clustering of coefficients for GA, BW, BL, and HC among the CpGs that yielded a Bonferroni-significant association for at least one birth outcome; overall, CpGs that were hypermethylated with exposure to MSDP were associated with reduced GA and birth size; while those that were hypomethylated with MSDP exposure were associated with increased GA and with increased birth size.

**Figure S9.** Correlation of regression coefficients of MSDP on DNAm from our meta-analyses of placental tissue vs. a prior meta-analysis of cord blood.

**Supplementary Methods:** Study specific data collection methods, funding information and acknowledgements.

**Supplementary Methods**

**AQUA (Asking QUestions about Alcohol in Pregnancy)**

Design and study population

The AQUA (Asking QUestions about Alcohol in Pregnancy) cohort was established to study the fetal effects of different patterns of prenatal alcohol exposure.^1^ Recruitment was conducted between July 2011 and July 2012 in several metropolitan public hospitals in Victoria, Australia. Eligibility criteria were women aged 16 years or more, who were less than 19 weeks in gestation, with an uncomplicated singleton pregnancy, and who were sufficiently proficient in English.

Maternal smoking during pregnancy (MSDP) definitions: any and sustained

Active maternal smoking during pregnancy was assessed via three self-administered questionnaires, which covered five time periods: three months prior to pregnancy, prior to pregnancy awareness, trimester one after pregnancy awareness, trimester two and trimester three.^1^ ‘Any maternal smoking’ was defined as “yes” if mothers reported to smoke at any time during pregnancy (even if smoking ceased after pregnancy recognition). For each prenatal period assessed, the number of cigarettes smoked per day was calculated as a continuous variable from a question on frequency (none, daily, weekly, fortnightly or monthly) and quantity per occasion (actual number).

Placenta collection and DNA extraction

Overall, 1570 participants were followed up until birth, and 248 placentas were collected. Three placental sections from each placenta were obtained randomly using a 6mm disposable biopsy punch. These biopsies were washed with sterile phosphate-buffered saline (PBS), dried, and then preserved in RNALater (Qiagen, Venlo, Netherlands) for 72 hours at 4˚C. The central segment of each biopsy was then dissected, and all three sections were pooled in a single vial, followed by storage at -20˚C until DNA extraction. Genomic DNA (gDNA) from 117 placental samples were extracted using the phenol/chloroform followed by ethanol precipitation method as previously described.^2^ Quality of gDNA was evaluated on a NanoDrop spectophotomer (Thermo Scientific, Waltham, MA, USA).

Placental genome-wide DNAm data acquisition, quality control and normalization

The extracted gDNA samples (500ng) were sent to Service XS (Leiden, The Netherlands) to generate DNAm data on the Illumina HumanMethylation450 (HM450) array. Samples were randomised and loaded onto ten arrays. Out of the 117 samples, 18 (15.4%) were excluded in this analysis because they were either of: non-white (i.e. not white Australian/UK/other European) ethnic origin (n=17) or had missing data for parity (n=1). Data was normalised based on Beta-Mixture Quantile (BMIQ) normalisation method and batch effect of 10 arrays was adjusted using ComBat Package in R.

Genome-wide differential DNAm analyses

Estimation of cell type proportions was performed using a reference free algorithm in R (RefFreeEWAS package), which estimated two cell types in our data. Robust linear regression models were used for genome-wide methylation analyses, adjusting for maternal age, parity, maternal education. For models that adjusted for cell types, only one cell type was included due to high correlation between the two estimated cell type proportions.

Birth outcomes

Birthweight and gestational age at birth were obtained from hospital records and maternal self-report. Birth length and head circumference at birth were not collected as part of this study.

Acknowledgements

The authors are extremely grateful to all the women and their children who took part in this study and wish to thank the researchers who were involved in sample collection, as well as Dr Sharon Lewis for providing the covariates for analysis.

Funding

This work was supported by the Australian National Health and Medical Research Council (Grant #1011070; Senior Research Fellowship #1021252 (JH) and the Victorian State Government’s Operational Infrastructure Support Program. The study has also received funding from the McCusker Charitable Trust to assist with the biospecimen collection at birth.

***EDEN (Study on the pre- and early postnatal determinants of child health and development)***

Design and study population

The Etude des Déterminants pré et post natals du développement et de la santé des Enfants (EDEN) is a population-based mother-child cohort study in France that aim to study the role pre- and post-natal factors in relation to child growth and development.^3^ Women were recruited between 2003 and 2006. More information about the EDEN cohort is available through our webpage <http://eden.vjf.inserm.fr/>. The EDEN cohort received approval from the ethics committee (CCPPRB) of Kremlin Bicêtre and from the French data privacy institution “Commission Nationale de l'Informatique et des Libertés” (CNIL). Written consent was obtained from the mother for herself and for the offspring.

Maternal smoking during pregnancy (MSDP) definitions: any and sustained

Active maternal smoking at each trimester of pregnancy was assessed using two questionnaires administered face-to-face to the mothers by trained interviewers between week 24 and 28 of pregnancy and at delivery. A-MSDP was defined as “yes” if mothers reported to smoke at any time during pregnancy and as “no” if mothers never smoked during pregnancy. S-MSDP was defined as “yes” when mothers reported to smoke at 1^st^ and 3^rd^ trimester of pregnancy or at 1^st^, 2^nd^ and 3^rd^ trimester of pregnancy; S-MSDP was defined as “no” if mothers never smoked during pregnancy or smoked during the 1^st^ and 2^nd^ trimester of pregnancy.

Placenta collection and DNA extraction

Overall, 1907 mother-child pairs were followed from until birth and placentas samples were collected for 1301 women. Placentas samples were collected at delivery by the midwife or the technician of the study using a standardized procedure. Samples of around 5mm x 5 mm were carried out in the centre of the placenta on the foetal side and were stored at −80 °C until processing. DNA from placental samples was extracted using the QIAsymphony instrument (Qiagen, Germany).

Placental genome-wide DNAm data acquisition, quality control and normalization

Genome-wide DNAm examination was performed in 668 placentas samples.^4^ The DNAm analysis was performed by the Centre National de Recherche en Génomique Humaine (CNRGH, Evry, France). The DNA samples were plated onto 96-well or 48-well plates. In total, nine plates including 64 chips were used. These plates were analyzed in 4 batches. The ratios for sex (boy/girl) and recruitment centre (Poitiers/Nancy) were balanced for each chip. Fifteen samples were measured in quadruplicates and one sample in duplicate across batches, sample plates and chips to detect technical issues such as batch effects. The Illumina's Infinium HumanMethylation450 BeadChip, representing over 485,000 individual CpG sites, was used to assess levels of methylation in placenta samples following the manufacturer's instructions (Illuminas, San Diego, CA, USA). Raw signals of 450 K BeadChips were extracted using the GenomeStudio® software (v2011.1. Illumina). The DNAm level of each CpG was calculated as the ratio of the intensity of fluorescent signals of the methylated alleles over the sum of methylated and unmethylated alleles (β value). All samples passed initial quality control and had on average>98% of valid data points (detection p-value < 0.01).

Genome-wide differential DNAm analyses

Genome-wide differential DNAm analyses were performed using robust linear regression models, adjusting for main covariates. Maternal age (years), parity (dichotomous), and maternal education (two-level factor) were obtained by questionnaires. In addition to the indicated covariates, models were also adjusted for center of recruitment. We used principal components analysis to assess batch-related variation and corrected for batch effects using comBat. Six components capturing cellular heterogeneity were considered for adjusting. Most participants were of white-European ethnic origin.

Birth outcomes

Birth weight and length were extracted from the maternity records. Gestational age was assessed from the date of the last menstrual period, and preterm birth defined as < 37 weeks of amenorrhea. Head circumference was the average of the two measurements performed by trainee midwifes during the postnatal examination of the neonate.

Acknowledgements

We thank the families that participated in the study for their generous contribution. We thank the midwife research assistants (L. Douhaud. S. Bedel. B. Lortholary. S. Gabriel. M. Rogeon. and M. Malinbaum) for data collection and P. Lavoine for checking, coding, and entering data. The EDEN mother-child cohort study group includes: I Annesi-Maesano, JY Bernard, J Botton, M-A Charles, P Dargent-Molina, B de Lauzon- Guillain, P Ducimetière, M de Agostini, B Foliguet, A Forhan, X Fritel, A Germa, V Goua, R Hankard, B Heude, M Kaminski, B Larroque, N Lelong, J Lepeule, G Magnin, L Marchand, C Nabet, F Pierre, R Slama, MJ Saurel-Cubizolles, M Schweitzer, O Thiebaugeorges.

Funding

This work was supported by the Fondation de France (n° 2012-00031593 and 2012-00031617), the French National Cancer Institute (INCa), the French Agency for National Research (ANR) (ANR-18-CE36-0005), the Fonds de Recherche en Santé Respiratoire, and the National Agency for Research (ANR-13-CESA-0011-02).

The EDEN cohort has been funded by the Foundation for Medical Research (FRM), National Agency for Research (ANR), National Institute for Research in Public Health (IRESP: TGIR cohorte santé 2008 program), French Ministry of Health (DGS), French Ministry of Research, Inserm Bone and Joint Diseases National Research (PRO-A) and Human Nutrition National Research Programs, Paris–Sud University, Nestlé, French National Institute for Population Health Surveillance (InVS), French National Institute for Health Education (INPES), the European Union FP7 programs (FP7/2007-2013, HELIX, ESCAPE, ENRIECO, Medall projects), Diabetes National Research Program (through a collaboration with the French Association of Diabetic Patients (AFD)), French Agency for Environmental Health Safety (now ANSES), Mutuelle Générale de l'Education Nationale (MGEN), French National Agency for Food Security, and the French-speaking association for the study of diabetes and metabolism (ALFEDIAM). Funders had no influence of any kind on analyses or interpretation of results.

***Gen3G (Genetics of Glucose regulation in Gestation and Growth)***

Design and study population

The Genetics of Glucose regulation in Gestation and Growth (Gen3G) is a prospective observational cohort study aiming to increase our understanding of biological, environmental, and genetic determinants of glucose regulation during pregnancy and their impact on foetal development and was described in details previously.^5^ In brief, we recruited a total of 1034 pregnant women aged ≥18 years old between January 2010 and June 2013 representing the general population of women in reproductive age receiving care at our institution. Women were excluded if they had non-singleton pregnancy, known pre-pregnancy diabetes or overt diabetes diagnosed based on biochemical screening that we performed at first trimester. The study protocol was approved by the Centre Hospitalier Universitaire de Sherbrooke (CHUS) ethic committee board and every participant gave written informed consent before enrolment in the study, in accordance with the Declaration of Helsinki.

Maternal smoking during pregnancy (MSDP) definitions: any and sustained

During the first research visit occurring at a mean of 9.4 weeks of gestation, trained study staff administered standardized questionnaires to participating women to collect medical history and lifestyle, including smoking behavior. Participants were classified as “any smoking” if reporting to smoke at the first visit or not currently smoking based on self-report at this visit. We did not collect data about smoking status later in pregnancy, thus we were unable to classify women as “sustained smoking”.

Placenta collection and DNA extraction

Placenta tissue from the fetal side was collected by trained study staff within 30-minutes of delivery. A 1-cm^3^ of placenta tissue sample was collected approximately 5-cm from the umbilical cord insertion. Placenta samples were subsequently stored at -80 ^0^C until DNA extraction occurred. We purified DNA from placenta samples using the All Prep DNA/RNA/Protein Mini Kit (Qiagen, USA). Purity of extracted DNA was evaluated using a Spectrophotometer (Ultrospec 2000 UV/Visible; Pharmacia Biotech, USA).

Placental genome-wide DNAm data acquisition, quality control and normalization

Within the Gen3G birth cohort we randomly selected 182 infants for the methylation study in the single pregnancies without gestational diabetes. From these 182 infants, we used DNA which was extracted from fetal placenta biopsies for the epigenome-wide DNAm analyses. To limit batch effects, we randomized all samples over the 96-well plates (including 6 duplicates). Samples (500 ng per sample) were placed on three 96-well plates. Bisulfite conversion was performed using the EZ-96 DNA Methylation kit (Zymo research Corporation, Irvine, USA). Then we used the Infinium HumanMethylation450 BeadChip (Illumina Inc., San Diego, USA) to measure the methylation level as a beta value ranging from 0 (no methylation) to 1 (complete methylation). During the quality control, we excluded 6 participants that were outliers in MDS plots. We performed DASEN normalization from the watermelon package. For this analyses, we excluded participants missing information exposure or covariates (n=10). We excluded probes that did not meet our criteria of a detection p-value<0.01 in more than 5% of the samples, and cross-reactive probes as indicated by Chen et al. The final analytic dataset included 166 samples and 398,741 probes

Genome-wide differential DNAm analyses

We conducted Epigenome-Wide-Association Analyses (EWAS) of self-reported prenatal maternal smoking behavior (any smoking) by fitting robust linear regression models using *MASS* package in R for each CpG on the normalized β-value scale adjusting for maternal age and parity. Maternal age was collected by questionnaire at first trimester visit. Parity was defined as the number of term pregnancies as asked in the first trimester questionnaire. We did not adjust for maternal social class and ancestry since data are not available in our study. All included participants were of European origin. In the second model, we further adjusted for putative cellular heterogeneity using the reference-free cell mixture decomposition method and we used 3 surrogate cell types.

Birth outcomes

Gestational age was obtained from the medical records: in clinic, gestational age is usually derived from last menstrual period (LMP), but when there was a discrepancy of more than 5 days between LMP and crown-rump length (CRL) estimated gestational age, CRL estimated gestational age was then used. Within two hours of delivery, birth weight (in g), birth length (in cm), and head circumference (in cm) were measured in the hospital following standard clinical protocol by obstetric nurses. Birth size z-scores were calculated using international references from the INTERGROWTH-21st Project.

Acknowledgements

Gen3G investigators acknowledge the Blood sampling in pregnancy clinic at the Centre Hospitalier de l'Universite de Sherbrooke (CHUS), and the assistance of clinical research nurses for recruiting women and obtaining consent for the study at the Research Center of CHUS. They also thank the CHUS Research in obstetrics services for organization of biosamples collection at delivery.

Funding

Gen3G was supported by Fonds de la recherche du Québec en santé (FRSQ) operation grant #20697 (M.F.H); a Canadian Institute of Health Research (CIHR) grant #MOP 115071 (M.F.H), and Diabète Québec grants (P.P. and L.B.). M.F.H. is supported by American Diabetes Association (ADA) Pathways to Stop Diabetes award (1-15-ACE-26).

***GENEIDA (Genetics, Early Life Environmental Exposures and Infant Development in Andalusia)***

Design and study population

The GENEIDA (Genetics, Early Life Environmental Exposures and Infant Development in Andalucía) project is a population-based prospective birth cohort of 802 mother-child pairs started in 2014 in the El Poniente (province of Almeria, South-Eastern Spain), this region is one of the most important intensive agricultural area in the country because of the large surface of plastic greenhouses. The aim of this project is to evaluate the role of early environmental exposures, such as chemical exposures, maternal stress, nutrition, on fetal and child grown and development. We are also studying the environmental influences on the placental epigenome and microbiome, and on maternal and children gut microbiome. The study was approved by the Research Ethics Committee of the province of Granada. The pregnant women received information of the study both written and orally, and they signed an informed consent.

Maternal smoking during pregnancy (MSDP) definitions: any and sustained

Active maternal smoking during pregnancy was assessed through a questionnaire administered face-to-face to the mothers by trained interviewers in the first (12-13 weeks) and third trimester (36 week) of pregnancy. Any maternal smoking was defined as “yes” if mothers reported to smoke at any time during pregnancy (even if they quit smoking before first trimester – week 12). Sustained maternal smoking was defined as “yes” when mothers reported to smoke at 1^st^ and 3^rd^ trimester of pregnancy.

Placenta collection and DNA extraction

Overall, 800 mother-child pairs were followed from until birth, a total of 631 placentas were collected. Each placenta, a 1cm^3^ biopsy was taken in 4 well-defined regions of chorionic villus at the fetal side and ~5cm far from the umbilical cord insertion. Biopsies were collected during the first two hours after birth. Samples were washed with cold 1X PBS in order to avoid maternal contamination. They were stocked in RNAlater at 4°C during 4 weeks at most. After that, RNAlater was removed and the samples were pulverized with liquid nitrogen and stocked in a biobank at -80°C until their processing.

DNA extraction procedures were carried out in 25 mg of placental tissue. Placental tissues were homogenizated 5 min to 50 Hz with stainless steel beads and 1mL of slagboom buffer using the TissueLyser LT (Qiagen). After the tissues is broken up, tissues were incubated with 20 uL of Proteinase K (20 mg/mL) at 37°C overnight, followed by digestion protocol proposed by Freeman et al. (2003)^6^ and modified by Gómez-Martín et al. (2015).^7^ DNA quality was evaluated on a NanoDrop 2000 spectrophotometer (Thermo Fisher Scientific, Waltham, MA). Isolated genomic DNA was stored at -80ºC until further processing

Placental genome-wide DNAm data acquisition, quality control and normalization

Genome-wide DNAm analysis was performed in 108 out of the 631 placental samples using the Infinium Human-Methylation450 BeadArray (Illumina, San Diego, CA USA) following manufacturer’s recommendations. All samples were randomly loaded onto the arrays and processed by the same technician at the same time to minimize batch effects and processed blind to sample identification at the GENYO, Centre for Genomics and Oncological Research: Pfizer/ University of Granada/ Andalusian Regional Government (Spain). DNA (500 ng) was bisulfite-converted with EZ DNA Methylation Kit (Zymo Research, Orange, CA) according to Illumina’s protocol for methylation profiling. BeadChips were scanned with an Illumina iScan software. Out of the 110 samples, 7 were excluded due to sex discordances (6.48%) and 14 because they were not of white-European ethnic origin (12.96 %) and 2 because information on maternal education was not available (1.85%).

Genome-wide differential DNAm analyses

Raw IDAT files were processed to extract methylation levels (beta-values) for each sample and normalized using Functional Normalization.^8^ Genome-wide differential DNAm analyses were performed using robust linear regression models, adjusting for main covariates of maternal age, parity, and maternal education, which were obtained by questionnaire administered to the mothers at first trimester of pregnancy. In addition, three components capturing cellular heterogeneity were considered for adjusting the model. All participants were of white-European ethnic origin. We abstracted birth weight measurements from medical registries

Birth outcomes

Gestational age (weeks), birth weight (grams), birth length (cm) and head circumference of the offspring were abstracted from medical records.

Acknowledgements

The GENEIDA researchers would like to thank all the mothers who have participated in this study, without whom this study would not have been possible. We would like also show our gratitude to medical and fieldwork staff of “Poniente Hospital” for their assistance in contacting the families, administering the questionnaires and collecting biological samples. Finally, we also thank lab staff from GENYO for their assistance in processing placental samples.

Funding

GENEIDA Project was funded by grants from Spanish Ministry of Health (FIS-PI13/01559) and Andalusian Ministry of Health (PI-0405-2014; PS-0205-2016; PS-0508-2016).

***Environment and Childhood Project (Infancia y Medio Ambiente, INMA)***

Design and study population

The Infancia y Medio Ambiente (INMA) (Environment and Childhood Project) is a population-based mother-child cohort study in Spain that aim to study the role of environmental pollutants in air, water and diet during pregnancy and early childhood in relation to child growth and development.^9^ More information about INMA project is available through our webpage http://www.proyectoinma.org/. Present study is based on the four *de novo* cohorts sited in Asturias, Gipuzkoa, Sabadell and Valencia and recruited between 2003 and 2008. The study was approved by the ethical committees of the centers involved in the study, and written informed consent was obtained from all the participants.

Maternal smoking during pregnancy (MSDP) definitions: any and sustained

Active maternal smoking during pregnancy was assessed through a questionnaire administered face-to-face to the mothers by trained interviewers in week 32 of pregnancy. A-MSDP was defined as “yes” if mothers reported to smoke at any time during pregnancy (even if they quit smoking before first trimester – week 12). S-MSDP was defined as “yes” when mothers reported to smoke at 1^st^ and 3^rd^ trimester of pregnancy.

Placenta collection and DNA extraction

Overall, 2506 mother-child pairs were followed from until birth and a random selection of 489 placentas were collected. Collected placentas at the four areas of study were stored at -80ºC in a central biobank until processing. Biopsies of approximately 5 cm^3^ were obtained from the inner region of the placenta, approximately 1.0-1.5 cm below the fetal membranes, corresponding to the villous parenchyma, and at a distance of ~5 cm from site of cord insertion. 25 mg of placental tissue were used for DNA extraction, previously rinsed twice during 5 minutes in 0.8mL of 0.5X PBS in order to remove traces of maternal blood. Genomic DNA from placenta was isolated using the DNAeasy® Blood and Tissue Kit, (Qiagen, CA, USA). DNA quality was evaluated on a NanoDrop spectophotomer (Thermo Scientific, Waltham, MA, USA) and additionally 100 ng of DNA were run on 1.3% agarose gels to confirm that samples did not present visual signs of degradation. Isolated genomic DNA was stored at -20ºC until further processing.

Placental genome-wide DNAm data acquisition, quality control and normalization

Genome-wide DNAm examination was performed in 190 out of the 489 placentas samples using the Infinium Human-Methylation450 array (Illumina, San Diego, CA USA) following manufacturer’s recommendations. All samples were randomly loaded onto the arrays and processed by the same technician at the same time to minimize batch effects and processed blind to sample identification at the University Medical Groningen Center UMCG Genome Analysis Facility (The Netherlands). DNA (500 ng) from each sample was treated by bisulfite conversion with the EZ-96 DNA Methylation Kit (Zymo Research) according to Illumina’s protocol for methylation profiling. BeadChips were scanned with an Illumina iScan and image data was uploaded into the Methylation Module of Illumina’s analysis software GenomeStudio (Illumina, San Diego, CA USA). Out of the 190 samples, 12 were excluded due to low performance (6.3%) and 10 due to sex discordances (5.3%).

Genome-wide differential DNAm analyses

Genome-wide differential DNAm analyses were performed using robust linear regression models, adjusting for main covariates. Maternal age, parity, and maternal education were obtained through a questionnaire administered to the mothers at first trimester. In addition to the indicated covariates, models were also adjusted for subcohort. The batch effect of 96-well bisulfite conversion plate (3 plates) was controlled using R ComBat Package (version 0.0.2). Two components capturing cellular heterogeneity were considered for adjusting. All participants were of white-European ethnic origin.

Birth outcomes

Birth weight and birth length of the offspring was recorded by trainee midwifes at delivery. We calculated gestational age from the date of the last menstrual period (LMP) reported at recruitment and confirmed using estimates based on the first ultrasound examination (about 12th week of gestation). When the difference between the LMP reported at recruitment and estimated from the ultrasound was ≥ 7 days, we estimated LMP using the crown-rump length.^10^

Acknowledgements

INMA researchers would like to thank all the participants for their generous collaboration. INMA researchers are grateful to Silvia Fochs, Nuria Pey, and Muriel Ferrer for their assistance in contacting the families and administering the questionnaires. The study was approved by the Ethical Committee of the Municipal Institute of Medical Investigation and by the Ethical Committee of the hospitals involved in the study. The pregnant women received information of the study both written and orally. Their informed consent of the participants was asked in each of the visits. A full roster of the INMA Project Investigators can be found at <http://www.proyectoinma.org/presentacion-inma/listado-investigadores/en_listado-investigadores.html>.

Funding

INMA was funded by grants from Instituto de Salud Carlos III (Red INMA G03/176, CB06/02/0041), Spanish Ministry of Health (FIS-PI041436, FIS-PI081151), Generalitat de Catalunya-CIRIT 1999SGR 00241, Fundació La marató de TV3 (090430). LAS was supported through a Colciencias PhD Scholarship, Colombia (Grant: 529/2011). CR-A was supported by a FI fellowship from Catalan Government (#016FI_B 00272).

***The New Hampshire Birth Cohort Study (NHBCS)***

Design and study population

The New Hampshire Birth Cohort Study (NHBCS) is an ongoing birth cohort that recruits pregnant women between the ages of 18 and 45 years, who are attending one of the study clinics in New Hampshire for prenatal care. This study was initiated to examine the impacts of environmental toxicants on growth and development, and thus only included those mothers who used an unregulated well as the primary source of drinking water. All participants provided written informed consent in accordance with the requirements of the Institutional Review Board (IRB) of Dartmouth College. Placenta (n=343) were sampled for genetic and epigenetic assays between February 2012 and September 2013. Interviewer administered questionnaires and medical record abstraction were utilized to collect sociodemographic, lifestyle, and anthropometric data.

Maternal smoking during pregnancy (MSDP) definitions: any and sustained

Maternal smoking during pregnancy (MSDP) was measured via a questionnaire which was administered via trained staff. Any MSDP was defined as self-reported smoking any number of cigarettes during any trimester of pregnancy.

Placenta collection and DNA extraction

Placental tissues were biopsied from the fetal side adjacent to the cord insertion site after removing maternal decidua and was performed within 2 hours of delivery. Samples were placed in RNAlater (Life Tecnologies, Carlsbad, CA) then frozen at -80°C. Both DNA was extracted (Norgen Biotek, Thorold, ON) and quantified via the Qubit Flourometer (Life Technologies), then subsequently stored at -80°C.

Placental genome-wide DNAm data acquisition, quality control and normalization

We measured DNAm via Illumina Infinium HumanMethylation450K BeadArray (Illumina, San Diego, CA) at the University of Minnesota Genomics Center. The EZ Methylation kit (Zymo Research, Irvine, CA) was utilized for bisulfite modification, samples were randomized across multiple batches, while batch variables were recorded. Data were assembled using BeadStudio (Illumina). The raw array data have been deposited at the NCBI Gene Expression Omnibus (GEO) for NHBCS via accession number GSE71678.

Genome-wide differential DNAm analyses

We used principal components analysis to assess batch-related variation and corrected for batch effects using comBat. Genome-wide differential DNAm analyses were performed using robust linear regression models, adjusting for maternal age (years), parity (dichotomous), and maternal education (three-level factor) which were measured via questionnaire. Two components capturing cellular heterogeneity were considered for adjusting. All participants were of white-European ethnic origin.

Birth outcomes

Gestational age was abstracted from medical records, and preterm birth defined as < 37 weeks gestation. Birth weight (grams), head circumference (cm), and birth length (cm), were abstracted from medical records.

Funding

This work was supported by the National Institutes of Health [NIH-NIGMS P20 GM104416, P01 ES022832]and by the United States Environmental Protection Agency [US EPA grant RD83544201]. Its contents are solely the responsibility of the grantee and do not necessarily represent the official views of the US EPA. Further, the US EPA does not endorse the purchase of any commercial products or services mentioned in the presentation.

***Rhode Island Child Health Study (RICHS)***

Design and study population

The Rhode Island Child Health Study (RICHS) is a study of mother-infant pairs with non-pathologic pregnancies that were enrolled from the Women and Infants’ Hospital in Providence, RI, USA between September 2010 and February 2013. Mothers younger than 18 years of age, with life threatening conditions, pregnancies resulting in preterm birth (< 37 weeks gestation), or congenital/chromosomal abnormalities were excluded. All protocols were approved by the institutional review boards at the Women and Infants Hospital of Rhode Island and Dartmouth College and all participants provided written informed consent. Infants born small for gestational age (≤ 10th BW percentile) or large for gestational age (≥ 90th BW percentile) were oversampled, then infants adequate for gestational age (between the 10th and 90th BW percentiles) that were matched on gestational age and maternal age were coincidentally enrolled. Sociodemographic and lifestyle data were collected via questionnaire while anthropometric and medical history data were obtained via structured medical record abstraction. Due to the vast majority of participants from other cohorts consisting of participants of white European ancestry, we restricted our analysis to those mothers that self-reported as white.

Maternal smoking during pregnancy (MSDP) definitions: any and sustained

Maternal smoking during pregnancy (MSDP) was measured via a questionnaire which was administered via trained staff. Those that reported smoking any cigarettes during any of the three trimesters were defined as those exposed to any MSDP.

Placenta collection and DNA extraction

Placental tissues were biopsied from the fetal side adjacent to the cord insertion site after removing maternal decidua and was performed within 2 hours of delivery. Samples were placed in RNAlater (Life Tecnologies, Carlsbad, CA) then frozen at -80°C. Both DNA was extracted (Norgen Biotek, Thorold, ON) and quantified via the Qubit Flourometer (Life Technologies), then subsequently stored at -80°C.

Placental genome-wide DNAm data acquisition, quality control and normalization

We measured DNAm via Illumina Infinium HumanMethylation450K BeadArray (Illumina, San Diego, CA) at the University of Minnesota Genomics Center. The EZ Methylation kit (Zymo Research, Irvine, CA) was utilized for bisulfite modification, samples were randomized across multiple batches, batch variables were recorded to allow for batch-effect corrections. Data were assembled using BeadStudio (Illumina). Raw array data are available via the NCBI Gene Expression Omnibus (GEO) for RICHS via accession number GSE75248. Principal components analysis was used to assess batch-related variations and corrected for batch effects using comBat.

Genome-wide differential DNAm analyses

We used principal components analysis to assess batch-related variation and corrected for batch effects using comBat. Genome-wide differential DNAm analyses were performed using robust linear regression models, adjusting for maternal age (years), parity (dichotomous), and maternal education (three-level factor) which were measured via questionnaire. Two components capturing cellular heterogeneity were considered for adjusting.

Placental gene expression measurement

Transcriptome-wide sequencing was performed on 194 placental samples. RNA was isolated with RNeasy Mini Kit (Qiagen, Valencia, CA), quantified via Nanodrop Spectrophotometer (Thermo Scientific, Waltham, MA), and and stored at -80°C. We assessed integrity via the Agilent Bioanalyzer (Agilent, Santa Clara, CA), removed ribosomal RNA via Ribo-Zero Kit,^11^ converted to cDNA using random hexamers (Thermo Scientific, Waltham, MA), and performed transcriptome-wide RNA sequencing via the HiSeq 2500 platform (Illumina, San Diego, CA).^12^ Raw reads are available at the NCBI sequence read archive (SRP095910). Quality control was performed in FastQC, then reads were mapped to the human reference genome (h19) using the Spliced Transcripts Alignment to a Reference (STAR) aligner. We excluded transcripts with very low expression levels, read counts were adjusted for GC content,^13^ then normalized via the trimmed mean of m-values (TMM);^14^ final data are normalized log2 counts per million (logCPM) reads. Because this was the only cohort that had both methylation and expression data, we aimed to include all available samples to have improved power to detect these associations. Thus this sample was from a racially heterogenous population, and we utilized EPISTRUCTURE^15^ to estimate and subsequently adjust for variability in the DNAm that may be attributed to ancestry or population structure.

Birth outcomes

Gestational age was abstracted from medical records, and there were no preterm births (< 37 weeks gestation) per the study exclusion criteria. Birth weight (grams), head circumference (cm), and birth length (cm), were abstracted from medical records.

Funding

This work was supported by the National Institutes of Health [NIH-NIMH R01MH094609, NIH-NIEHS R01ES022223].

**Funding information for SJL**

SJ London is supported by the Intramural Research Program of the NIH, National Institute of Environmental Health Sciences (ZO1 ES49019). This work was also supported by the National Institutes of Health [NIH-NIMH R01MH094609, NIH-NIEHS R01ES022223 and the Intramural Research Program of the NIH-NIEHS, ESZ0149019].
