## Supplemental Figures for "Placental DNA methylation signatures of maternal smoking during pregnancy and potential impacts on fetal growth"

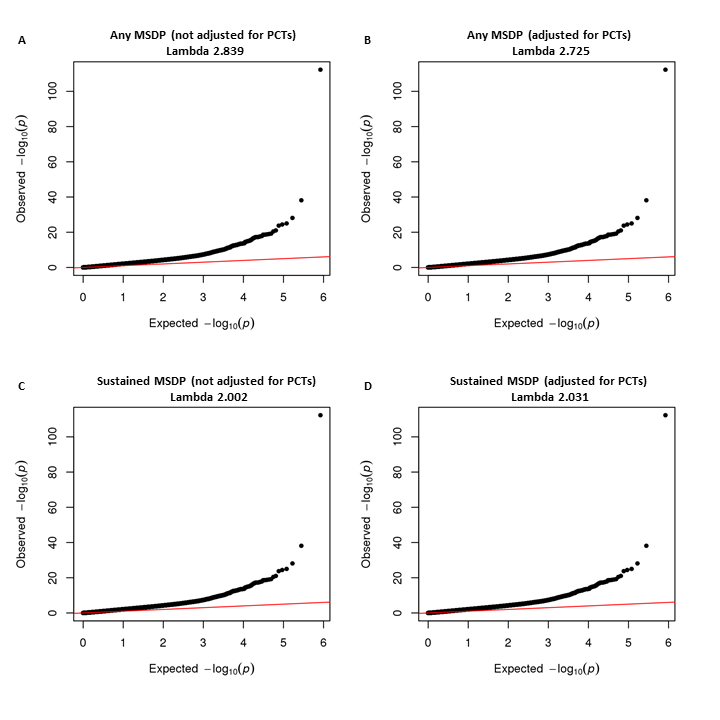

**Figure S1.** QQ-plot for the meta-analyses of the association between placental DNAm and any MSDP without (A) and with adjustment for cellular heterogeneity (B) and sustained MSDP without (C) and with adjustment for cellular heterogeneity (D); N=1700 (A and B) and N=795 (C and D); PCT = putative cell type proportions.

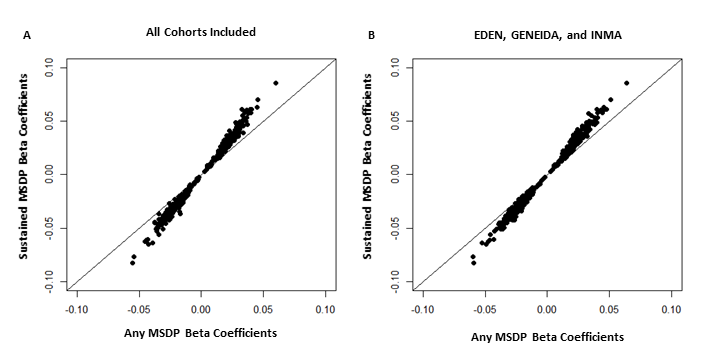

**Figure S2.** Plots showing the increase in the magnitude of coefficients for models of sustained MSDP versus any MSDP, (A) when the meta-analysis included all particiapting cohorts, and (B) when meta-analysis only included those cohorts that also participated in models of sustained MSDP (EDEN, GENEIDA and INMA); this analysis included the 548 CpGs that were Bonferroni-sgnificant for associations with any and sustained MSDP. One CpG (cg27402634) was excluded from plot due to being an outlier and making visualization of the differences difficult for the associations with any and sustained MSDP; this site also showed a larger magnitude of effect in the model when all cohorts were included (% difference in the size of the beta-coefficient between sustained and any MSDP = 12.7%), and when only EDEN, GENEIDA, and INMA were included (% difference in the size of the beta-coefficient between sustained and any MSDP = 27.5%).

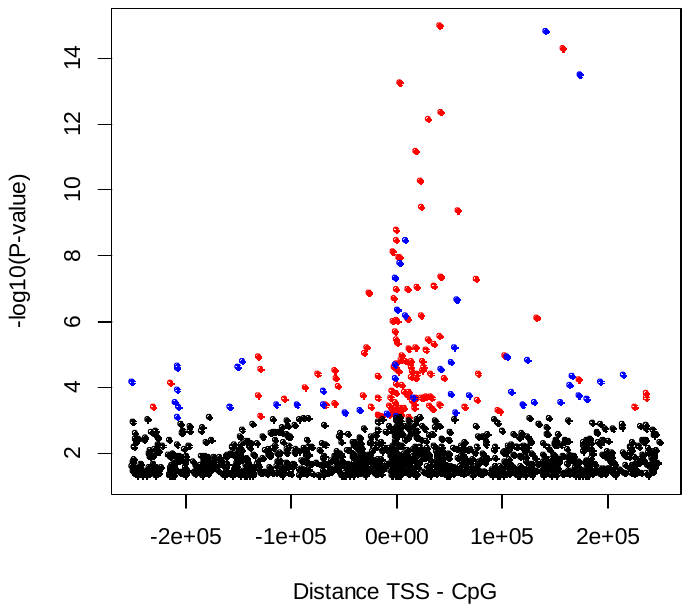

**Figure S3**. –log10**(***P*) versus distance between gene transcription start site (TSS) and CpG site for 1219 eQTM in which DNAm was positively associated with expression at 5% FDR (blue), or inversely associated with expression at 5% FDR (red).

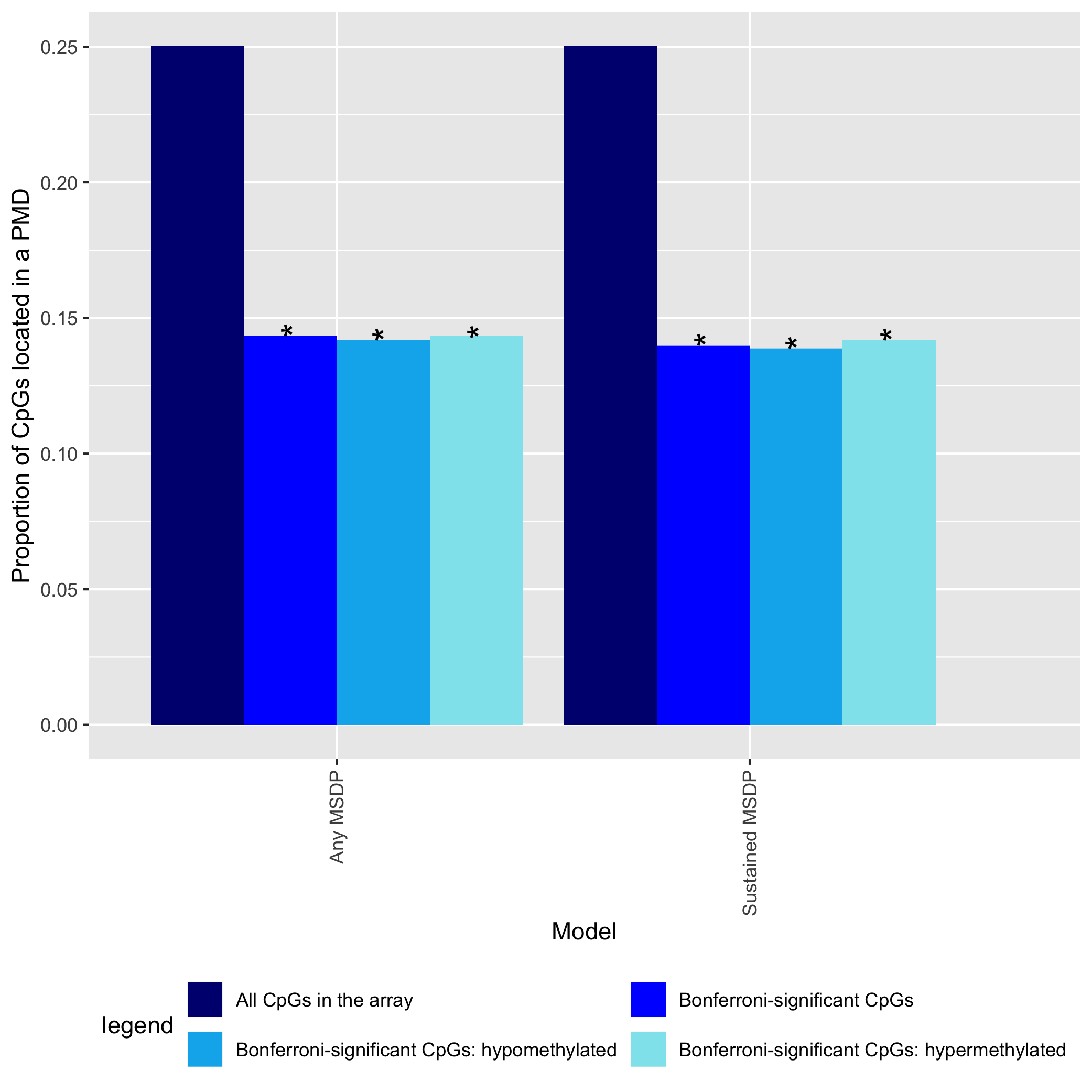

**Figure S4.** Proportion of CpGs annotated to placental partially methylated domains (PMDs), by significance and direction of the effect for associations with any and sustained MSDP.

*Nominally significant P-value via the hypergeometric test

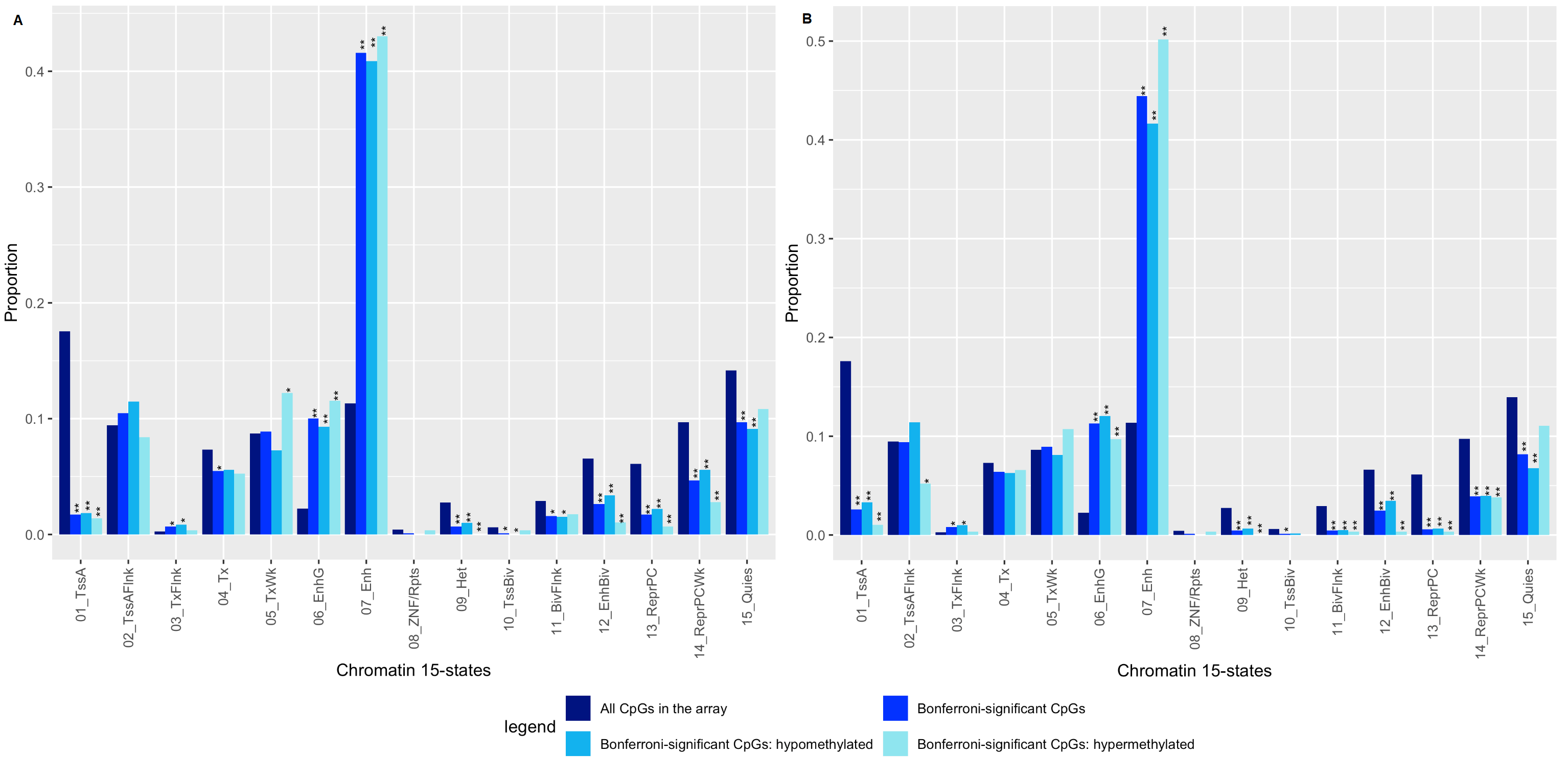

**Figure S5.** Proportion of CpGs annotated to each of the 15 placenta chromatin states from ROADMAP, by significance and direction of the effect for associations with any MSDP (A) and sustained MSDP (B).

Chromatin states are defined as: 01_TssA = Active transcription start site; 02_TssAFlnk = Flanking active transcription start site; 03_TxFlnk = transcribed state at the 5’ and 3’ end of genes showing both promoter and enhancer signatures; 04_Tx = Actively transcribed state (strong transcription); 05_TxWk = Actively transcribed state (weak transcription); 06_EnhG = Genic enhancer; 07_Enh = Enhancer; 08_ZNF/Rpts = state associated with zinc finger protein genes and repeats; 09_Het = Consitutive heterochromatin; 10_TssBiv = Bivalent/Poised TSS; 11_BivFlnk = Flanking bivalent TSS/Enh; 12_EnhBiv = Bivalent enhancer; 13_ReprPC = repressed polycomb; 14_ReprPCWk = Weak repressed polycomb; 15_Quies = Quiescent state.

*Nominally significant P-value in the hypergeometric test

**Bonferroni significant P-value (p-value< 0.003)

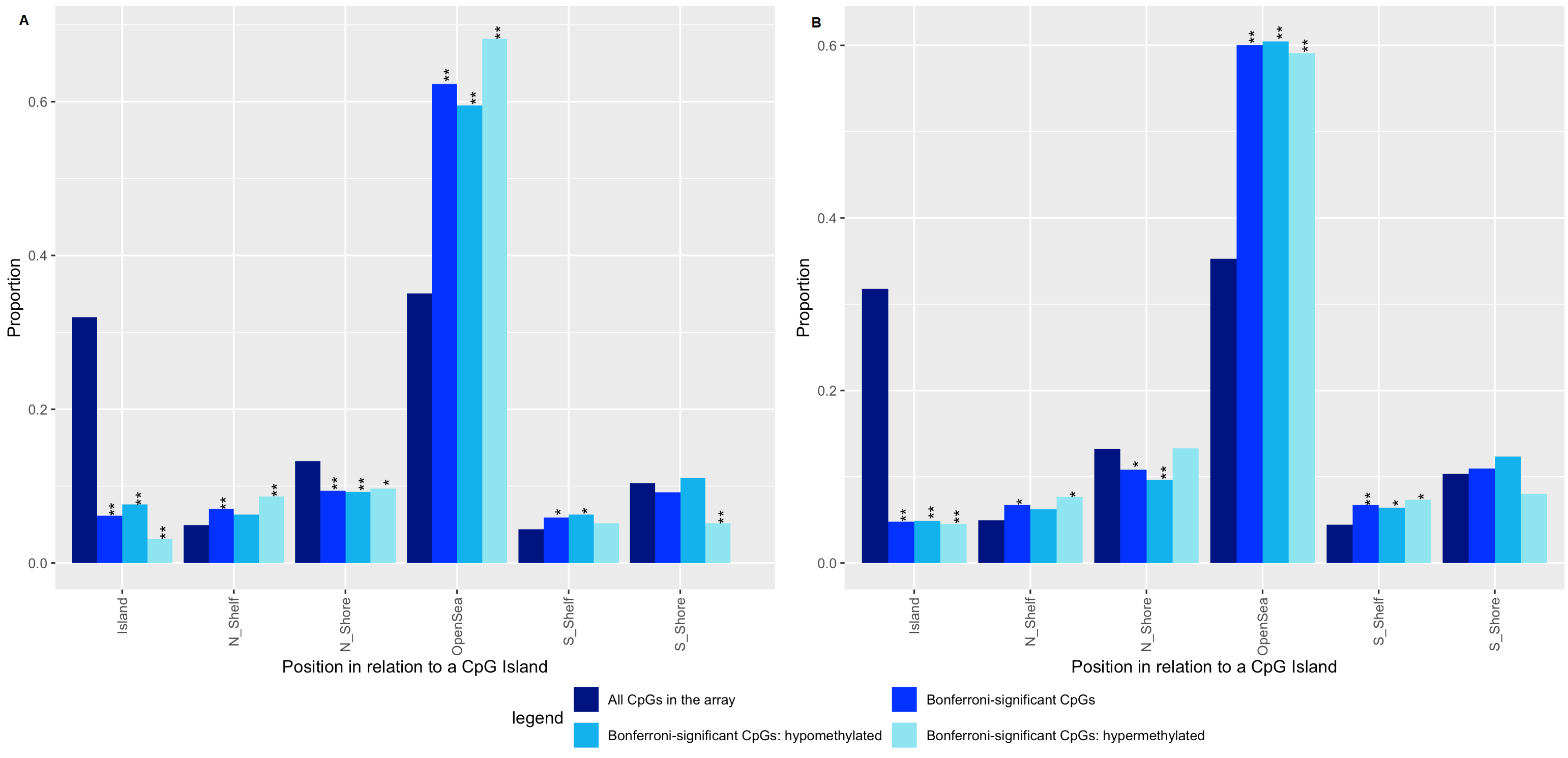

**Figure S6.** Proportion of CpGs annotated to each of the 6 locations in relation to a CpG island, by significance and direction of the effect, for those CpGs associated with any (A) and sustained (B) MSDP.

*Nominally significant P-value in the hypergeometric test

**Bonferroni significant P-value (p-value<0.008)

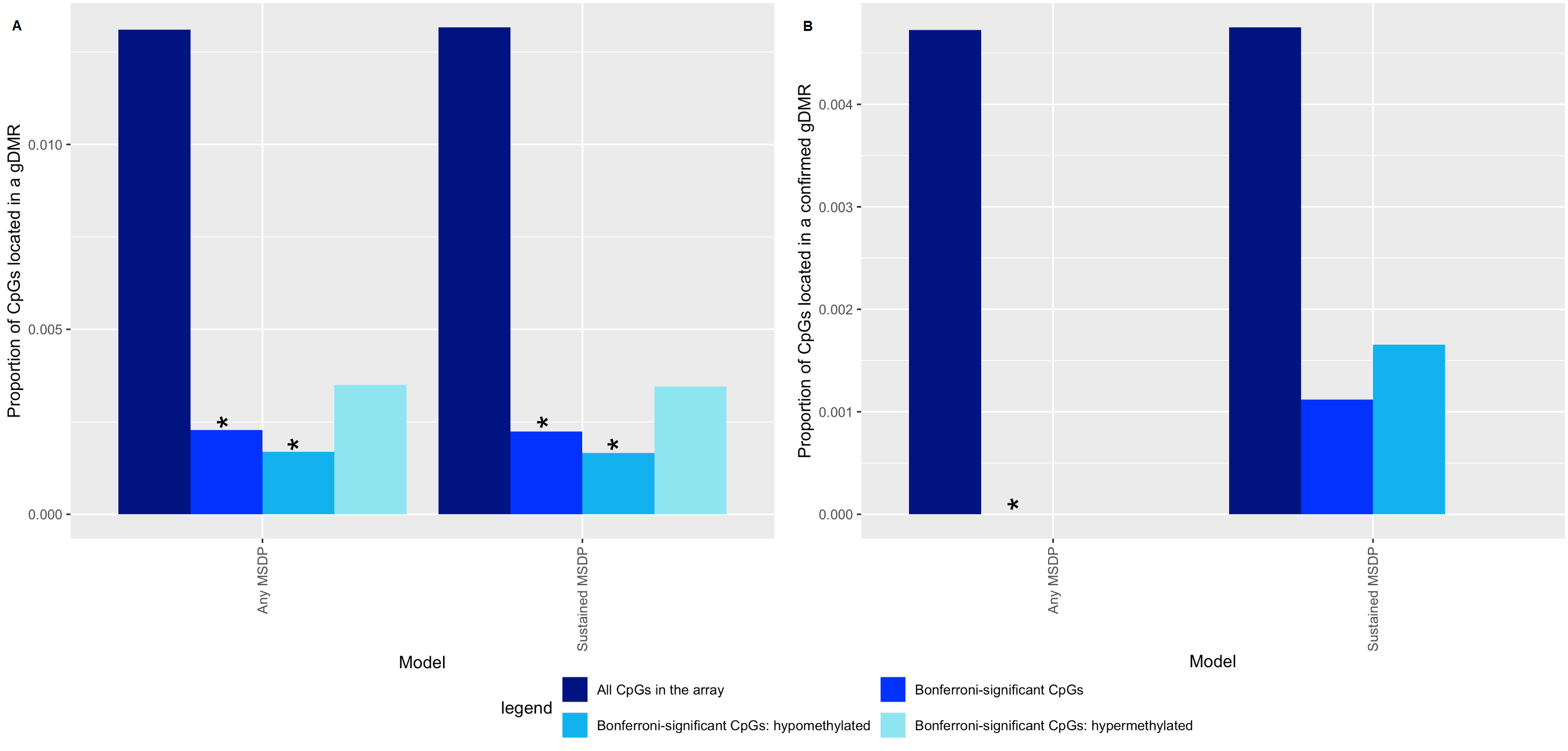

**Figure S7.** Proportion of CpGs annotated to candidate (A) or confirmed (B) placental germline differentially methylated regions (gDMR), by significance and direction of the effect for those CpGs associated with any and sustained MSDP.

*Nominally significant P-value in the hypergeometric test

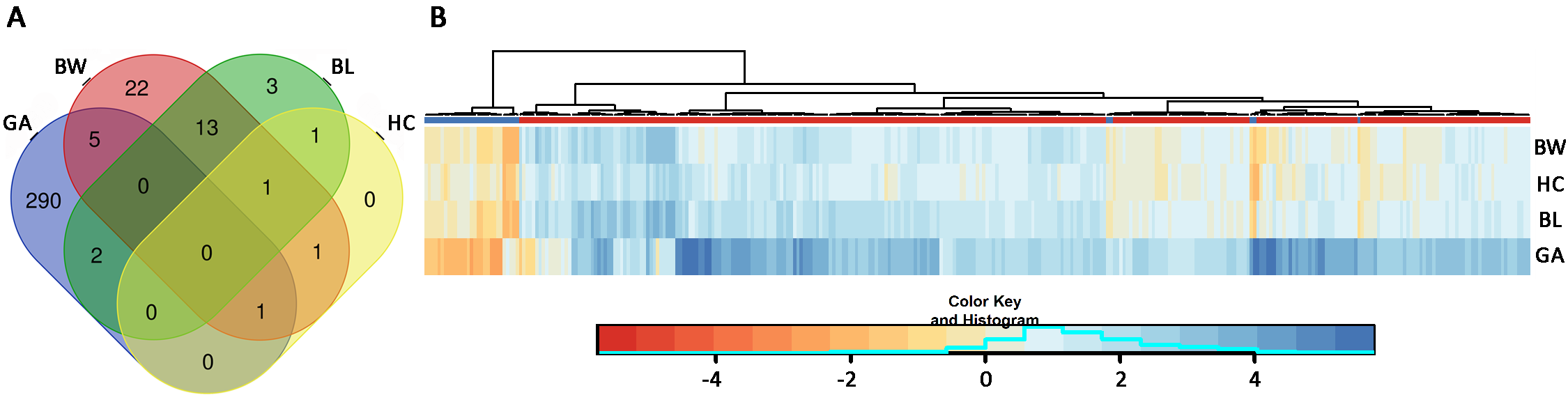

**Figure S8.** Associations between DNAm at 1224 smoking-sensitive CpGs and birth outcomes. (A) Venn diagram of the 339 CpGs that were associated with gestational age (GA), birth weight (BW), birth length (BL) and head circumference (HC) at Bonferroni significance (1224 CpGs tested), and (B) Clustering of coefficients for GA, BW, BL, and HC among the CpGs that yielded a Bonferroni-significant association for at least one birth outcome (blue = positive, red = inverse); top axis color bar shows direction of association between DNAm and MSDP (blue = positive, red = inverse) from the meta-analysis results. In general, CpGs that were hypermethylated with exposure to MSDP were associated with reduced GA and birth size; while those that were hypomethylated with MSDP exposure were associated with increased GA and with increased birth size.

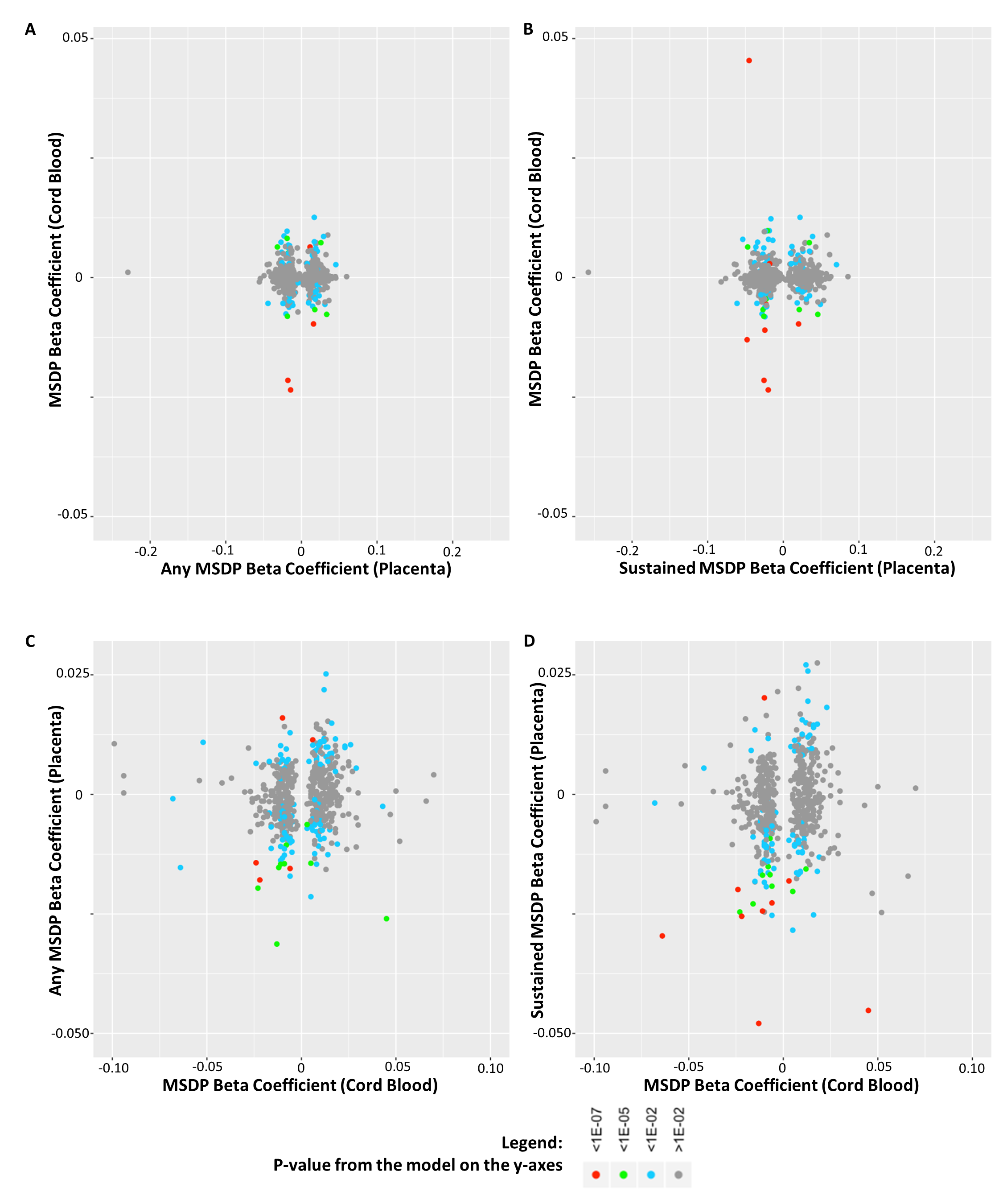

**Figure S9.** Correlation of regression coefficients of MSDP on DNAm in placenta vs. cord blood. The coefficients for CpGs that yielded associations within a 5% FDR for a particular tissue are presented on the x-axis, while the coefficients for those CpGs in the other tissue are presented on the y-axis; raw meta-analysis p-values for the models on the y-axis are indicated with different colors (see legend). We observed no correlation between cord blood and placental differential methylation for CpGs that were associated (5% FDR) with any (A; r^2^=0.04) or sustained (B; r^2^=0.04) MSDP in our meta-analyses; nor did we observe any correlation in differential methylation between cord blood and placenta for the CpGs that were associated with MSDP in the prior meta-analysis of cord blood (C; r^2^=0.09 and D; r^2^=0.08). Cord blood results were retrieved from Joubert et al. (2016) for sustained MSDP adjusted for cell type heterogeneity.
